# XLID Syndrome Gene Med12 Promotes Ig Isotype Switching through Chromatin Modification and Enhancer RNA regulation

**DOI:** 10.1101/2022.06.14.496024

**Authors:** Farazul Haque, Tasuku Honjo, Nasim A. Begum

## Abstract

The transcriptional co-activator Med12 regulates gene expression through the function of its kinase module and by interacting with the larger Mediator complex and associated RNA Polymerase II (RNAPII). Here, we show a kinase module-independent function of Med12 in antibody class switching recombination (CSR). Med12 is essential for IgH 3’ regulatory region (3’RR) or super-enhancer activation and functions with p300 and Jmjd6/Carm1 coactivator complexes. Med12 deficiency leads to a dramatic decrease in H3K27 acetylation and enhancer RNA (eRNA) transcription at 3’RR, with concomitant impairment of AID-induced DNA double strand breaks, long-range S-S synapse formation, and 3’RR-Eμ interaction. CRISPR/dCas9-mediated enhancer activation re-establishes the epigenomic and transcriptional hallmarks of the 3’RR super-enhancer, fully restoring Med12 depletion defects. Notably, we find that 3’RR derived eRNAs are critical for promoting proper S region epigenetic regulation, S-S synapse formation and recruitment of Med12 and AID to the IgH locus. We find specific X-Linked intellectual disability syndrome associated Med12 mutations are defective in both 3’RR eRNA transcription and CSR, suggesting B and neuronal cells may have cell-specific super-enhancer dysfunctions. We conclude Med12 is essential for IgH3’RR activation and eRNA transcription and plays a central role in AID-induced antibody gene diversification and genomic instability in B cells.

## INTRODUCTION

Upon antigen encounter, mature B cells undergo immunoglobulin (Ig) class switch recombination (CSR) to allow transition of Ig isotype expression from IgM to IgG, IgA or IgE(*1*). At the genetic level, the IgM constant gene is replaced by one of the downstream constant genes through Ig locus specific DNA rearrangement induced by activation-induced deaminase (AID)(*2, 3*). This programmed DNA arrangement involves 3 critical steps: (1) germline transcription through long intronic S regions, (2) targeted DNA breaks in recombining S regions, and (3) joining cleaved S region pairs by NHEJ-mediated repair(*4–9*). AID plays a central role in these processes by inducing DNA cleavage at specific loci and promoting S-S synapse (unique *cis* conformation) formation to facilitate recombination(*10, 11*). Although the requirement of AID in DNA cleavage and S-S synapse formation is well established, the precise mechanism of locus specificity and S-S tethering have yet to be fully understood.

The IgH 3’RR super enhancer region plays an important role in AID-induced CSR(*12, 13*) and somatic hypermutation (SHM)(*14*), but not in RAG-induced V(D)J recombination(*15*). However, how 3’RR *cis*-regulatory elements influence and/or regulate AID-induced actions during CSR remains poorly understood. The most prevalent and widely-accepted view suggests the 3’RR is mainly involved in germline transcription (GLT) (*13, 16, 17*), which is one of the essential prerequisites that precedes AID-induced CSR. The core 3’RR encompasses four transcriptional enhancers (hs3a, hs1.2, hs3b and hs4) and deletion of all four decreases the GLTs, with expected decreases in CSR(*13, 17*). These 3’RR enhancers are thought to function co-operatively, as their individual deletion does not impair Ig isotype switching. However, deletion of both hs3b and hs4 decreases CSR for all isotypes except IgG(*12*). In summary, the precise mechanisms underlying 3’RR dependent CSR regulation remains unsettled, despite studies demonstrating the cumulative activity of individual enhancers, such as hs1.2 and hs4, or in combination with hs3b(*18*).

Enhancers and super-enhancers are typically considered to be distally located gene regulatory elements that enhance transcription by long-distance interaction with gene *promoters*(*19–21*). Enhancer activation requires formation of specific protein complexes that include transcription factors (TFs), coregulators, chromatin remodelers and modifiers, and RNAPII(*19–21*). One enhancer coactivator is the supramolecular Mediator complex, which serves as a signaling coordinator between transcriptional machinery and RNAPII through modulating subunit composition(*22, 23*). Mediator is assembled as four subcomplexes referred to as “head”, “middle” “tail” and “kinase” modules(*24–26*). While the head, middle and tail modules comprise the core Mediator, the fourth module reversibly interacts with the core through its Med13 subunit. The kinase module is composed of Med12, Med13, Cdk8, and Cyclin C and contributes to both negative and positive regulation of RNAPII-driven transcription(*26–29*).

The N-terminal L domain of Med12 is responsible for CDK8 activation, representing a kinase module-dependent function of Med12(*30*). Notably, while Leukemia and Uterine Leiomyoma associated mutations are localized within the L domain, X-linked intellectual disability (XLID) associated mutations, including atypical and typical FG, Lujan, Opitz-Kaveggia or Ohdo syndrome, are distributed across the central LS domain(*31, 32*). Several prostate cancer related mutations are also located in the LS and PQL domain(*33, 34*). The PQL domain is adjacent to the LS domain and is subject to arginine methylation at several positions, which likely contributes to ncRNA binding and protein-protein interactions(*35*). Additionally, some PQL domain mutations may confer chemotherapeutic resistance in cancer(*36*). Mutations in the C-terminal OPA domain of MED12 has been reported in three siblings with low IgG levels and B-cell proliferation defects(*37*). Although Med12 mutations mentioned are thought to be crucial for disease progression, the functional significance and molecular mechanisms of these mutations remain poorly understood.

Here, we use various domain-specific Med12 mutations to determine Med12 structure-function relationships within the context of AID induced genomic instability in mature B cells. We find Med12 is dispensable for switch germline transcription but is indispensable for 3’RR activation and transcription. We also find these Med12 functions are independent of kinase module function. Mechanistically, Med12, in collaboration with p300 and Jmjd6/Carm1 complexes, preserves signature histone epigenetic marks at 3’RR and S regions, resulting in adequate eRNA production and unperturbed CSR. Loss of Med12 leads to both architectural and epigenomic dysregulation of the IgH locus, abrogating AID-induced DNA double strand breaks and post-break recombination/repair processes essential for Ig isotype switching.

## RESULTS

### Med12 is required for AID-induced DNA breaks and S-S synapse formation during CSR

To investigate the requirement of Med12 in CSR, we established an siRNA-mediated Med12 gene knockdown (Med12^KD^) model in the murine B cell line CH12F3-2A. This cell line efficiently switches from IgM to IgA in response to stimulation with CD40L, IL-4, and TGFβ (CIT)(*38*). Multiple siMed12 concentrations effectively inhibited CSR with a clear trend of dose dependency (Fig. 1a and Extended Data Fig 1a). We subsequently used 20pMol of siMed12 in all the experiments as it did not affect cell proliferation. To confirm knockdown specificity, we designed a wild-type Med12 construct (WT^R^) capable of producing Med12 transcripts resistant to degradation by siMed12 (Fig. 1b, Top). Transfection of cells with the Med12 WT^R^ construct showed a dose dependent CSR complementation in Med12^KD^ cells (Fig. 1b). These results suggest Med12 plays a critical role in CSR and may be specifically required for effective CSR.

**Fig. 1.**
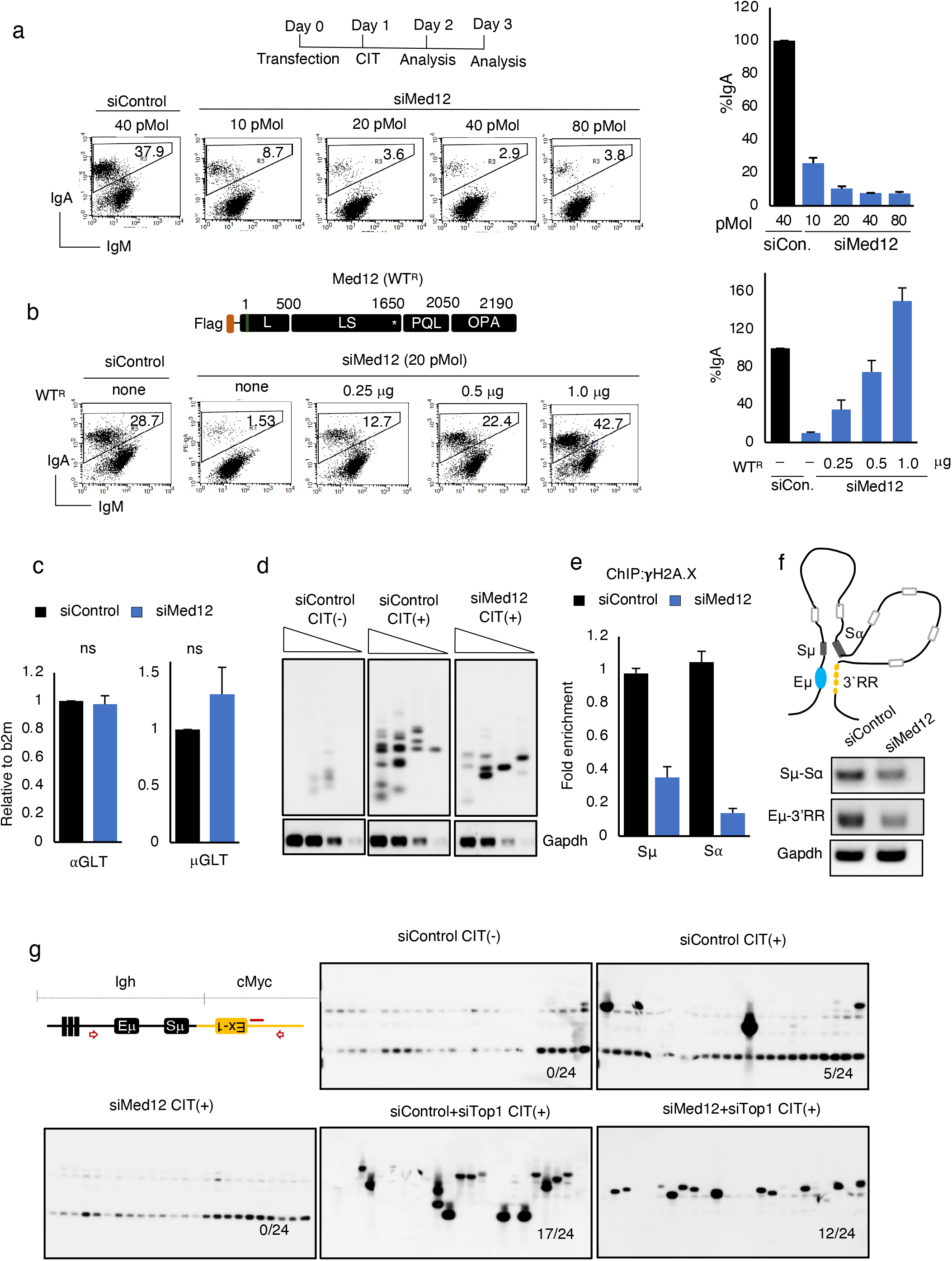
Med12 depletion leads to impaired AID-induced DNA break, S-S synapse and *Igh/c-Myc* translocation. **a,** The flowchart showing the time course of the experiment. The CH12F3-2A cells was transfected with either siControl or siMed12 (Med12^KD^) and stimulated (+) by CIT as indicated and analyzed by Flow cytometry analysis (FACS). (Bottom) The FACS analysis showing the effect of different doses of siMed12 as indicated on IgA switching compared to siControl. (Right) the bar graph showing the mean values of IgA switching from three independent experiments and the data represents the mean ± sd, statistical significance was calculated by two-tailed Student’s t-test (P*<0.05). **b,** Representation of siMed12 resistance wild-type Med12^R^ (WT^R^) used for the CSR rescue experiment, asterisk (*) mark shows the site of the si-resistance. The WT^R^ construct tagged with 3xFlag at their N-terminal and the lengths of each domain of Med12 protein showed by the amino acid’s positions. The FACS analysis showing the effect of different concentrations of WT^R^ on IgA complementation efficiencies after 30h CIT (+) in siControl and Med12^KD^ CH12F3-2A cells. (Right) the bar graph showing the mean values of IgA switching and the data represents the mean ± sd calculated from three independent experiments. Statistical significance was calculated by two-tailed Student’s t-test (P*<0.05). **c,** The RT-qPCR bar graph showing the effect of siControl (black bar) and siMed12 (20pMol) (blue bar) on aGLT and μGLT transcripts. The value has been normalized with endogenous beta-2 microglobulin (b2m) abundance. The result summarizes the means ± s.d. of three independents experiments and the statistical significance was determined by two-tailed Student’s t-test (P>0.05), ns indicates in significant difference. **d,** LM-PCR based DNA break assay showing the amplified Sμ region in CIT (-) and CIT (+) (siControl/siMed12) CH12F3-2A cells. The samples treated with siControl or siMed12 were collected and processed followed by southern blot with the 5’-Sμ specific probe. Bottom panel shows the semiquantitative PCR analysis of Gapdh of respected samples as an internal control. The triangles indicate a threefold dilution in the DNA amount. The blot represents the best of three independent experiments. **e,** The ChIP-qPCR assay for γH2AX occupancy at S regions (Igh locus) in siControl and siMed12 treated CH12F3-2A cells. Values were normalized to the DNA input signals followed by the maximum value in each data set. The results summarize the means ± s.d. of three independents experiments and the statistical significance was determined by two-tailed Student’s t-test (P*<0.05). **f,** (Up) Schematic view of the long-range interactions occurred at the Igh locus during CIT induced IgA switching in CH12F3-2A cells, which brings Sμ and Sα into close proximity. (Bottom) Representative gel picture of the chromosome confirmation capture PCR (3C-PCR) analysis, detecting the long-range interactions between different Igh regions in siControl and siMed12 treated CH12F3-2A cells. Each PCR panel shows the combinations of primer pairs used for recombined DNA amplification. Gapdh PCR analysis of the cross-linked DNA sample served as a loading control. **g,** (Left) PCR amplification scheme to detect Igh/c-Myc chromosomal translocations. Red triangles represent the positions of nested PCR primers used to amplify the rearranged regions. The position of the Myc probe used in the southern blot hybridization is shown as a horizontal red bar. (Rght) Southern blot analysis of PCR-amplified fragments with a Myc-specific probe from two independent experiments. CH12F3-2A cells were transfected with the indicated siRNAs and stimulated the cells for 48h followed by DNA isolation and PCR.

We confirmed that Med12 deficiency did not affect switch germline transcription (GLTs), μGLT and aGLT, or AID expression (Fig. 1c and Extended Data Fig. 1a). Consistently, Med12 depletion did not affect AID-ER (AID fused with Estrogen Receptor-ER) expression but strongly impaired CSR in AIDER-CH12F3-2A line (Extended Data Fig 1b,c). To examine the exact steps of CSR that are impaired following Med12 depletion, we assessed AID-induced DNA double strand break (DSB) formation in the S region by linker ligation-mediated PCR (LM-PCR)(*39*). Med12 depletion results in a sharp decrease in DNA double strand break signals, which are restored to control levels with co-transfection of the Med12 WT^R^ construct (Fig. 1d, Extended Data Fig 1d), suggesting AID-induced DNA double strand break formation requires Med12. Consistently, gH2AX, a marker of early DNA damage response (DDR) signaling, was also diminished at the Sμ and Sα regions in Med12^KD^ cells (Fig. 1e).

IgA CSR requires a unique *cis* conformation of the IgH locus known as the S-S synapse. This synapse brings the donor Sμ and the acceptor Sα into close proximity by looping out a large intervening sequence region. S-S synapse formation detection assays have included standard chromosome confirmation capture (3C), which enables detection of S-S proximity as a hybrid PCR product of two widely separated loci, namely Sμ and Sα. Similarly, 3C assays can be used to detect the proximity or long-range interactions between promoters and enhancers (Eμ-3’RR) or interaction with novel loci other than IgH. In Med12^KD^ cells, we find the intensity of Sμ-Sα hybrid PCR products was strongly decreased, (Fig. 1f), suggesting Med12 is required for proper S-S synapse formation during CSR. Depletion of Med12 also markedly diminished the interaction between Eμ and 3’RR, providing further evidence for the role of Med12 for both constitutive (Eμ-3’RR) and inducible (S-S) locus conformation.

As AID expression induces IgH/c-Myc translocation in addition to CSR in B lymphocytes, we next examined IgH/c-Myc translocation frequency by PCR amplifying translocated genomic DNA junctions from siControl and siMed12 transfected cells. PCR products were subjected to Southern hybridization using c-Myc locus-specific probe(*40, 41*). Med12 depletion resulted in a reduction of AID-induced IgH/c-Myc translocations (Fig. 1g). Similarly, Med12 depletion also counteracted augmentation of IgH/c-Myc translocations by Top1 deficiency(*42*).

Ligation of DSB ends between two S regions mainly occurs through the nonhomologous end joining (NHEJ)–mediated DNA repair pathway, we next examined the impact of Med12 deficiency in a NHEJ mediated repair assay using *I-SceI* meganuclease induced DSBs(*43*). In this assay, a GFP reporter gene is expressed only after successful NHEJ between two cleaved *I-SceI* sites, removing an intervening thymidine kinase (TK) gene cassette. The *I-Sce*I-expressing plasmid was co-transfected with siControl or siMed12 into the reporter cell line, H1299dA3-1. As expected, in siControl treated cells, *I-SceI* expression showed 12.6% GFP (+) cells, which was reduced to 3.4% in siMed12 treated cells, suggesting DSB end-joining via NHEJ requires Med12 (Extended Data Fig 1e). Taken together, we conclude Med12 is vital for promoting AID-induced DNA double strand breaks, S-S synapse formation, and NHEJ - all of which are essential for Ig isotype switching.

### S region DNA break and synapse formation are regulated by distinct domains of Med12

To understand the molecular basis of the diverse functions of Med12, we examined Med12 structure-function relationships by generating several deletion and point mutant constructs. Med12 is a large protein with four distinct domains (Fig. 2a): a leucine-rich domain (L), a leucine and serine rich domain (LS), a proline, glutamine and leucine rich domain (PQL), and an odd paired area glutamine rich domain (OPA). We generated individual domain specific deletions using WT^R^ sequence as a template (Fig. 2a). CSR complementation efficiency of these mutants was measured relative to Med12^KD^ cells transfected with the WT^R^ construct. We find the ΔL^R^, ΔOPA^R^, and ΔNLS^R^ mutants display wild-type level CSR complementation, whereas ΔLS^R^ and ΔPQL^R^ show a significant loss of CSR activity (Fig. 2b). Interestingly, we find that when CSR defective ΔLS^R^ and ΔPQL^R^ constructs are co-transfected, CSR is restored back to WT^R^ levels (Fig. 2b). However, as individual LS and PQL domains were intact in the respective ΔPQL^R^ and ΔLS^R^ mutants, we suspect functional cross-complementation occurs between these two domains.

**Fig. 2.**
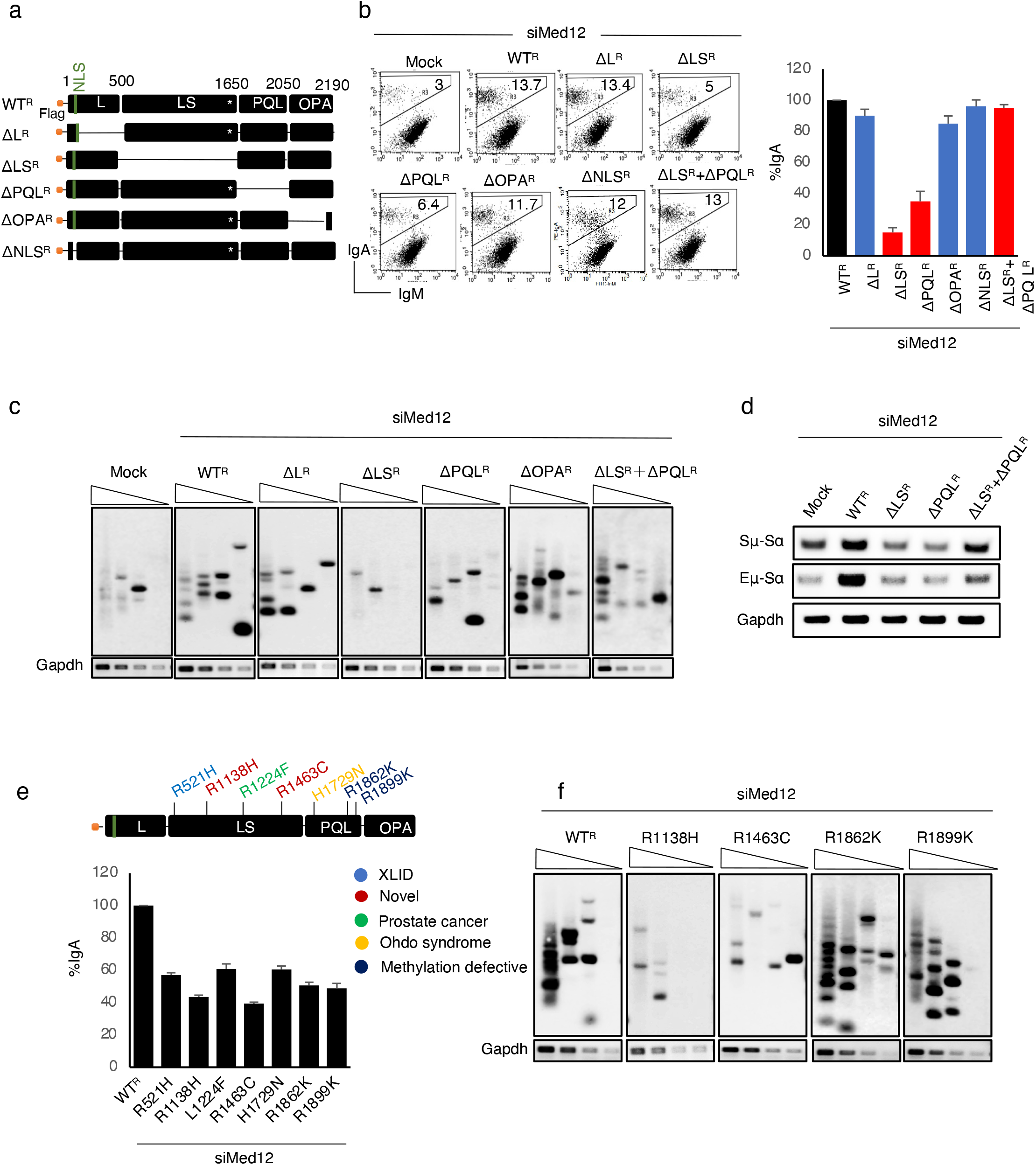
Med12 regulates CSR differentially by its LS and PQL domains. **a,** Schematic representation of various Med12 deletion mutants used in the IgA complementation experiment. NLS, nuclear localization signal. **b,** FACS analysis showing the IgA rescue efficiency by transiently transfected WTR and various Med12^R^ deletion mutants in Med12^KD^ CH12F3-2A cells. The samples have been analyzed after 24h of CIT (+). Mock contains empty backbone vector. Right, the bar graph summarizes the means ± s.d. of five independents experiments and the statistical significance was determined by two-tailed Student’s t-test (P*<0.05). **c,** LM-PCR essay to detect the AID induced DNA break in transiently transfected WT^R^ and corresponding domain deleted Med12^R^ constructs in Med12^KD^ CH12F3-2A cells followed by southern blot with the 5’-Sμ specific probe. The samples have been collected after 30h of CIT (+) and proceed similarly as described in fig 1d. Gapdh used as internal control. The triangles indicate a threefold dilution in the DNA amount. **d,** 3C-PCR analysis in WT^R^ and indicated Med12 deletion constructs in Med12^KD^ CH12F3-2A cells. The samples have been collected after 48h of CIT (+) and processed. Each PCR panel shows the combinations of primer pairs used for DNA amplification. Gapdh PCR of the cross-linked DNA sample served as a loading control. **e,** (Top) Schematic representation of the WT^R^ Med12 protein and its associated various disease linked point mutations scattered over LS and PQL domains. The different color code corresponds to linked disease. (Bottom) The bar plot showing the IgA rescue efficiency in disease linked Med12 mutations including the two novel mutations found during the study in Med12^KD^ CH12F3-2A cells. Graph summarizes the means ±s.d. of three independents experiments and the statistical significance was determined by two-tailed Student’s t-test (P*<0.05).. **f,** LM-PCR essay for estimation AID induced DNA break rescue in WT^R^ and corresponding disease linked-CSR defective mutants in Med12^KD^ CH12F3-2A cells. Gapdh used as internal control. The triangles indicate a threefold dilution in the DNA amount.

To further examine this possible cross-complementation, we tested whether ΔLS^R^ and ΔPQL^R^ mutants can support AID-induced DNA double strand breaks and S-S synapse formation by LM-PCR and 3C assays. LM-PCR results suggest the ΔLS^R^ mutant is severely defective in S region DSB formation in Med12^KD^ cells, which is consistent with this mutant’s CSR deficiency (Fig. 2b). Conversely, S region DNA break signals were detected in Med12^KD^ cells transfected with ΔL^R^, ΔPQL^R^, ΔLS^R^+ΔPQL^R^, and ΔOPA^R^ mutants (Fig. 2c). Notably, ΔPQL^R^ rescued the severe DNA break defect in ΔLS^R^ transfected Med12^KD^ cells. Taken together, we conclude the LS domain of Med12 plays a critical role in AID-induced S-region DNA break.

We next examined the contribution of the ΔLS^R^ and ΔPQL^R^ mutants and their co-expression (ΔLS^R^+ΔPQL^R^) in S-S synapse formation. We find Med12 depletion significantly reduces Sμ-Sα and Eμ-Sα interactions, while WT^R^ transfection of Med12^KD^ cells restores these interactions (Fig. 2d). We find neither ΔLS^R^ nor ΔPQL^R^ transfection restores Sμ-Sα or Eμ-Sα interaction in Med12^KD^ cells. However, we find that co-transfection of both ΔLS^R^ and ΔPQL^R^ rescues Sμ-Sα and Eμ-Sα interactions, suggesting both the LS and PQL domain are required for synapse formation.

To examine whether Med12 mutations observed in human disease have CSR defects, we tested several reported point mutations across multiple Med12 protein domains (Extended Data Table1). Since LS domain is associated with XLID, we also included two novel LS domain mutations (R1138H, R1463C) that were generated during the site-directed mutagenesis process. We find mutations in the LS (R521H, R1138H, L1224F, and R1463C) and PQL (H1729N, R1862K, and R1899K) domains show diminished CSR activity (Fig. 2e, Extended Data Table 1). Point mutants that induced significant CSR defects (R1138H, R1463C, R1862K, and R1899K) were further examined for their ability to promote AID-induced DNA double strand breaks. Remarkably, LM-PCR assays show that impaired DNA double strand break formation in Med12^KD^ cells can be restored by complementation with PQL domain mutants (R1862K, R1899K), but not by LS domain mutants (R1138H, R1463C) (Fig. 2f). Despite promoting DNA double strand breaks as efficiently as WT Med12, PQL mutants were CSR defective (Fig. 2e, Extended Data Table1), likely due to their recombination defects. Taken together, we suggest Med12 regulates two distinct steps of CSR, namely AID-induced DNA double strand breaks using its LS domain and S region synapse formation using its PQL domain.

### The transcription regulatory kinase module of Med12 is dispensable for CSR

The N-terminal L domain of Med12 plays an important role in transcriptional regulation through its interaction with the kinase module and the core Mediator(*24*). The Mediator kinase module is comprised of Med12, Cyclin C, Cdk8/Cdk19, and Med13. This kinase module reversibly interacts with the core Mediator through Med13 and contributes to gene expression regulation(*44*) (Fig. 3a). As Med12 stimulates Cdk8/19 kinase activity through Cyclin C (CycC) dependent interaction, loss of Med12 or a defect at the N-terminus disrupts kinase complex formation, resulting in gene expression defects(*44*). To test the requirement of the kinase module and the core Mediator for CSR, we treated CH12F3-2A cells by Cortistatin A (CA), a potent pharmacological inhibitor of Cdk8 and its paralog Cdk19(*45, 46*). Surprisingly, CA treatment (100 nM or 250 nM) did not show any inhibitory effect on CSR compared to DMSO control (Fig. 3b, Extended Data Fig. 2a). We confirmed the effectiveness of CA treatment (250 nM) by examining expression of kinase module regulated and CA-sensitive genes, namely: Egr1, Atf1, and Atf7(*47*) (Fig. 3c, Extended Data Fig. 2b). Additionally, the depletion of Cdk8 by siRNA had no significant effect on CSR (Fig. 3d, Extended Data Fig. 2c), despite Cdk8 mRNA being efficiently downregulated by three independent siCdk8s (Fig. 3e, Extended Data Fig. 2c).

**Fig. 3.**
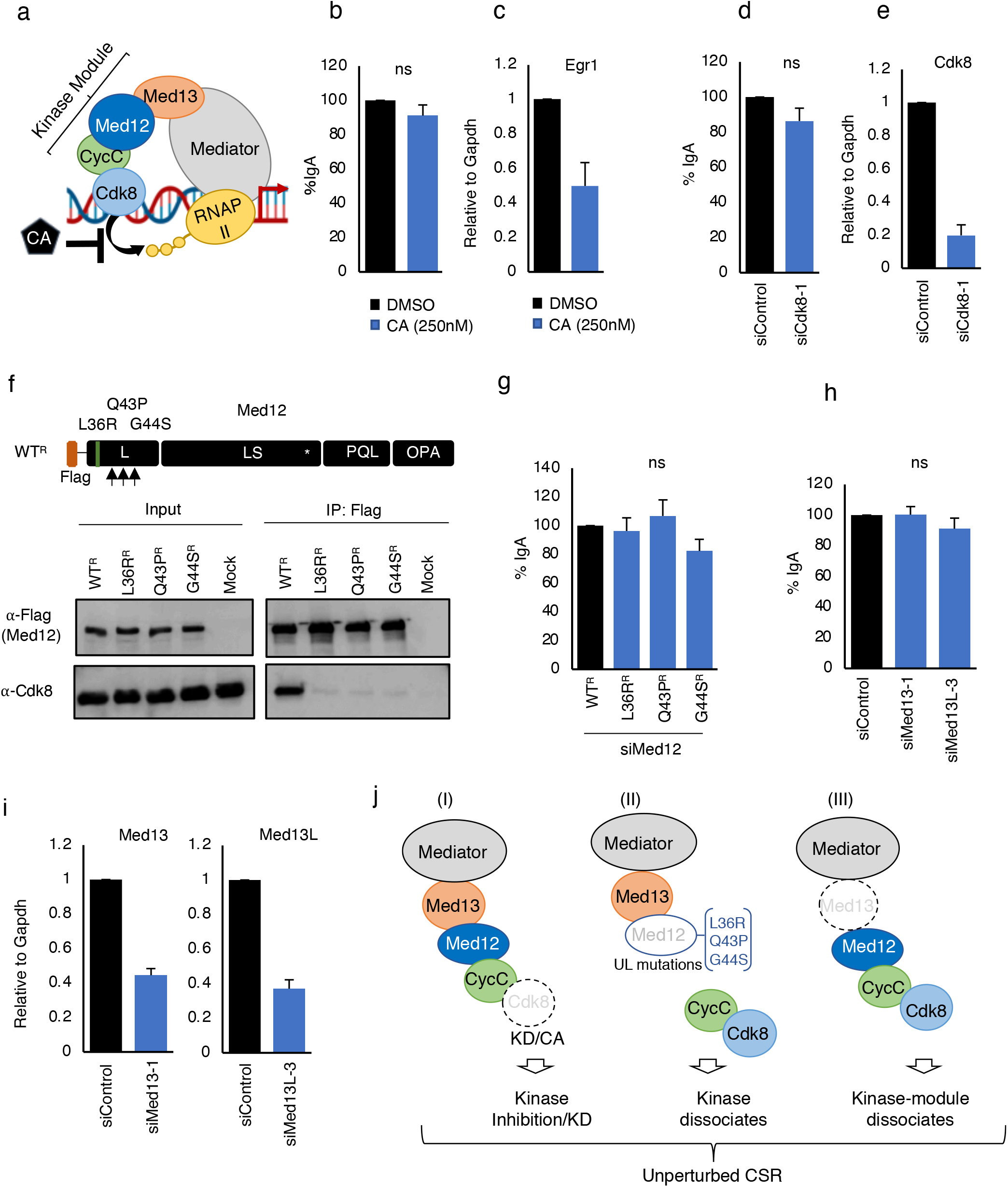
Med12 does not require kinase and core Mediator complex for CSR. **a,** Schematic representation depicting the Med12 kinase module and core Mediator function involved in transcription regulation. Med12 kinase module consist of four proteins (Med12, Med13, CcyC and cdk8) and Med13 act as an anchor and connects the kinase module to the core Mediator complex. Cortistatin A (CA) inhibits the Med12 kinase activity by binding to Cdk8 and inhibits the RNAPII phosphorylation and gene transcription. **b,** The representative bar plot showing the effect of CA on IgA switching from three independent experiments. The CIT (+) stimulated CH12F3-2A cells were treated with control (DMSO) or CA (250nM) for 24h and later IgA efficiency was estimated by FACS analysis. **c**, The cells were collected for RNA isolation followed by cDNA synthesis. Egr1 transcripts abundance was estimated by RT-qPCR in DMSO and CA treated cells and used as positive control. The experiments were performed on three independent days and the statistical significance was determined by two-tailed Student’s t-test. ns-non significance (P*<0.05). **d,** The bar plot showing the effect of Cdk8 knockdown (KD) on IgA switching and **e,** corresponding RT-qPCR analysis to determine the KD efficiency by siCdk8. The values have been normalized to endogenous Gapdh. ns-non significance (P>0.05). **f,** (Up) Schematic representation of Uterine leiomyoma linked point mutations found on Med12.The arrow indicates the location of the mutagenesis. (Bottom) The HEK293T cells were transfected by lipofectamine either with indicated Med12 constructs for 48h and later samples were collected for overnight Flag immunoprecipitation and later processed for gel chromatography. Input shows (10%) of the IP samples and the blot was developed against the indicated antibodies. Mock shows untransfected cells. **g,** The bar plot showing the effect of IgA complementation by Uterine leiomyoma linked Med12 point mutations in Med12^KD^ CH12F3-2A cells. The experiments were performed on three independent days and the statistical significance was determined by two-tailed Student’s t-test. ns-non significance (P>0.05). **h, i** The effect of Med13 and Med13L knockdown on IgA switching. The CH12F3-2A cells were transfected with either siMed13 or siMed13L and CIT (+) stimulated for 24h and analyzed by FACS. The samples have been collected and processed for RT-qPCR analysis to determine the KD efficiency by each siRNAs. The values have been normalized to endogenous Gapdh. The experiments were performed on three independent days and the statistical significance was determined by two-tailed Student’s t-test. ns-non significance (P>0.05). **j,** The schematic showing the step wise dissection of the kinase domain and there components requirement for CSR. (I) Cdk8 initiates the kinase activity and phosphorylates the RNA polII for gene transcription.CA or Cdk8 depletion inhibits the kinase activity. (II) The Uterine leiomyoma linked Med12 point mutations (L36R, Q43P,G44S) dissociates the CycC/Cdk8 complex from the kinase domain although Med12/Med13 remain attached to the core Mediator.(III) Med13/Med13L links kinase domain to core Mediator whereas depletion of Med13/Med13L dissociates Med12-CycC-Cdk8 from core Mediator, in all the cases CSR has been unchanged.

To further investigate kinase module independent functions of Med12 in CSR, we modeled three Med12 mutations (L36R, Q43P and G44S) that are associated with Uterine Leiomyoma(*48*) (Fig. 3f). These mutations disrupt Med12 interaction with Cyclin C and Cdk8/Cdk19. Notably, each single mutation alone was sufficient to abolish the kinase module dependent functions of Med12(*48*). We confirmed loss of Cdk8 interaction in these Med12 mutants by co-immunoprecipitation (Co-IP) (Fig. 3f). Furthermore, all three mutants fully restored CSR activity. (Fig. 3g). These results are consistent with the insensitivity of CSR to CA treatment and Cdk8 depletion (Fig. 3b,d) and further suggest kinase module independent functions of Med12 in CSR (Fig. 3j).

Furthermore, to test the requirement of core Mediator in CSR, we knocked down Med13 and its paralogue Med13Like (Med13L) which is required for kinase module interaction with core Mediator (Fig. 3a). Med13 and Med13L were depleted up to 70-80% by siMed13-1 and siMed13L-3, respectively. However, CSR was not significantly affected following Med13 or Med13L depletion (Fig. 3h,i and Extended Data Fig. 2d,e). We observed a slight decrease in CSR by siMed13-2/3 that we deemed as nonspecific, as it did not correlate with knockdown efficiency (Extended Data Fig. 2d). Taken together, we conclude from these multiple lines of evidence that Med12 promotes CSR independently of its kinase module and the core Mediator (Fig. 3j).

### Med12-p300 complex regulates CSR through 3’RR enhancer activation

Previous reports have largely focused on the kinase module-mediated functions of Med12, leaving the kinase module or core Mediator independent functions of Med12 largely unknown. A recent report suggests Med12 regulates hematopoietic stem cell (HSC) specific super-enhancers in co-operation with the histone acetyltransferase (HAT) p300 and independent of the Med12 kinase module(*27*). From this, we asked whether Med12 might analogously activate the IgH 3’RR.

To test this hypothesis, we first examined whether deposition of the hallmark enhancer marker, H3K27ac, at the IgH 3’RR super-enhancer was dependent on Med12 and p300. We find that depletion of either Med12 or p300 dramatically reduces H3K27ac deposition across the enhancer clusters (hs3a, hs1.2, hs3b, hs4, hs5, hs6, and hs7) in 3’RR (Fig. 4 a-c, Extended Data Fig. 3a). Since p300 is a key acetyltransferase that catalyzes H3K27 acetylation at enhancers, we tested whether p300 recruitment at the IgH 3’RR was dependent on Med12. Indeed, Med12 depletion significantly reduced p300 localization at the 3’RR enhancer (Fig. 4d). In support of this finding, co-IP analysis confirms the interaction between Med12 and p300 (Extended Data Fig. 3b). Furthermore, functional depletion of p300 either by siRNA or HAT catalytic inhibition by C646 inhibitor (p300^In^), led to CSR impairment (Fig. 4e, Extended Data Fig. 3a,c) and decreased H3K27ac at the enhancer.

**Fig. 4.**
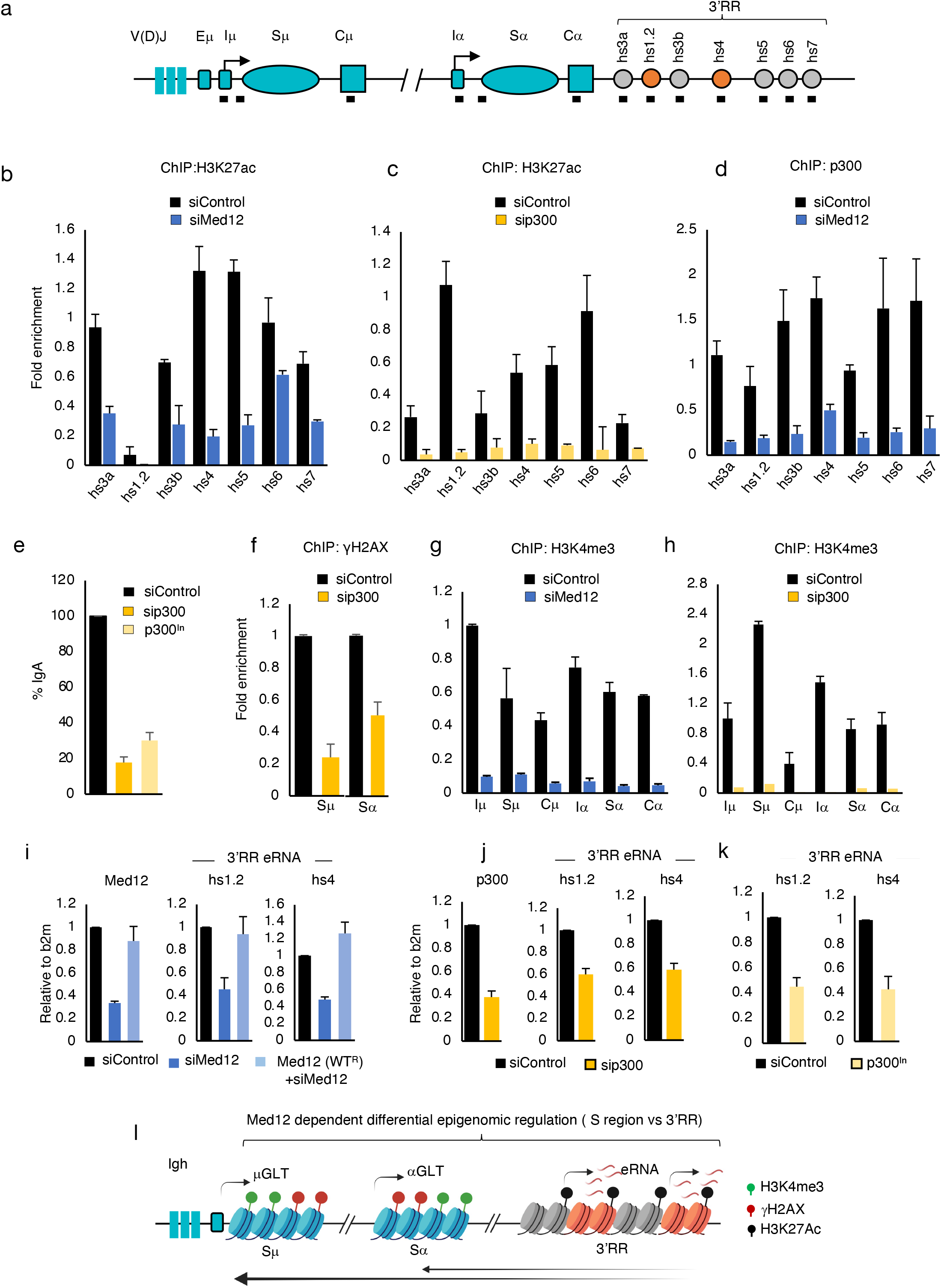
3’RR enhancer activation by Med12-p300 complex is required for AID-induced DNA break and S-S synapse. **a,** Schematic showing the different regions present at the Igh locus, the black bar showing the position of the primers used for ChIP-qPCR amplification. **b, c,d,** The ChIP-qPCR showing the enrichment in control and Med12 or p300 knockdown cells using the indicated antibodies. The values were normalized to the DNA input signals followed by the maximum value in each data set. The experiments were performed on two independent days and the statistical significance was determined by twotailed Student’s t-test (P*<0.05). **e,** The bar plot showing p300 knockdown by siRNA and inhibition of p300 HAT (p300^In^) activity by C646 (5nM) and there effect on CSR. The samples were transfected and analyzed as mentioned before. **f, g,h** The ChIP-qPCR showing the relative enrichment in control and knockdown cells as indicated. The antibody used for ChIP is indicated on each panel. The values were normalized to the DNA input signals followed by the maximum value in each data set. The experiments were performed on two independent days and the statistical significance was determined by two-tailed Student’s t-test (P*<0.05). **I,j,k,** The RT-qPCR data showing the effect of Med12 and p300 knockdown or p300^In^ on indicated transcripts relative to control. The data was normalized with endogenous b2m abundance. **l,** Schematic showing the role of Med12 dependent differential epigenomic regulation at S and 3’RR regions. We propose that Med12 activates the 3’RR enhancer through p300 which marks H3 histone acetylation (black boll). The eRNA produced form the activated enhancers travelled far from the enhancers and regulates the AID-induced DNA break formation by regulating the histone marks gH2AX (Red boll) and H3K4me3 (Green boll) at S regions.

To confirm p300 loss indeed recapitulates the Med12 loss, we investigated the three essential steps for CSR: 1) switch germline transcription, 2) DNA double strand break formation, and 3) S-S synapse formation. Surprisingly, p300 depletion decreased S region double strand breaks as well as interactions between Sμ-Sα and Eμ-3’RR (Extended Data Fig. 3e,f,g), similar to the defects observed following Med12 depletion (Fig. 1d,f). Also similar to Med12^KD^ (Fig. 1c,), transcription of μGLT and αGLT were not perturbed following p300 depletion (Extended Data Fig. 3h). However, p300 knockdown or its catalytic inhibition decreased DSB associated gH2AX formation in the S region (Fig. 4f, Extended Data Fig. 3d). Given this observation, we examined H3K4me3 marks in the S region, as they are essential for AID-induced DNA double strand *breaks*(*49–52*). Remarkably, depletion of Med12 or p300 resulted in a significant loss of H3K4me3 at the S region (Fig. 4g,h). We observed a similar loss of H3K4me3 from both the Sμ and Sα regions following p300 inhibition (Extended Data Fig. 3d). Co-IP experiments also showed that the Med12-p300 complex contains critical components involved in histone lysine-4 methylation, such as Ash2 and Wdr5 (Extended Data Fig. 3b). Loss of either Ash2 or Wdr5 also significantly reduced S region H3K4me3 and gH2AX formation, resulting in impaired AID induced DSB and CSR(*50*).

In summary, 3’RR activation through the coordinated activity of Med12 and p300 appears to be crucial for both S region chromatin remodeling and long-range S-S synapse formation. To further investigate this hypothesis, we tested whether transcription of noncoding enhancer RNAs (eRNA), another hallmark of active enhancers, was also dependent on Med12 and/or p300. We find that eRNA transcription from hs1.2 and hs4 is decreased by Med12 depletion and elevated by Med12 overexpression in Med12^KD^ cells (Fig. 4i). Similarly, p300 depletion or HAT activity inhibition reduces 3’RR eRNA expression (Fig. 4j-k), suggesting 3’RR enhancer activation is dependent on Med12 and/or p300. We conclude that Med12 plays a crucial role in recruiting p300 to the 3’RR for enhancer activation and epigenomic modulation in the S region, both of which are essential for efficient CSR (Fig. 4l).

### Rescue of Med12 deficiency by CRISPR/dCas9 mediated 3’RR activation

We hypothesized that the loss of 3’RR activation is the underlying cause of CSR impairment following Med12 deficiency. From this, we tested whether CSR can be restored by enforced enhancer activation using a CRISPR/dCas9 based enhancer-epigenetic remodeling system(*53, 54*). We employed a hs1.2 and/or hs4 targeting dCas9 coupled to the core HAT domain of p300 (dCas9-p300^C^) that can catalyze H3K27 acetylation at the 3’RR enhancer(*55*) (Fig. 5a). We selected the two hypersensitive sites, hs1.2 and hs4, for targeting in CH12F3-2A as their deletion significantly impairs CSR in mouse models(*56, 57*).

**Fig. 5.**
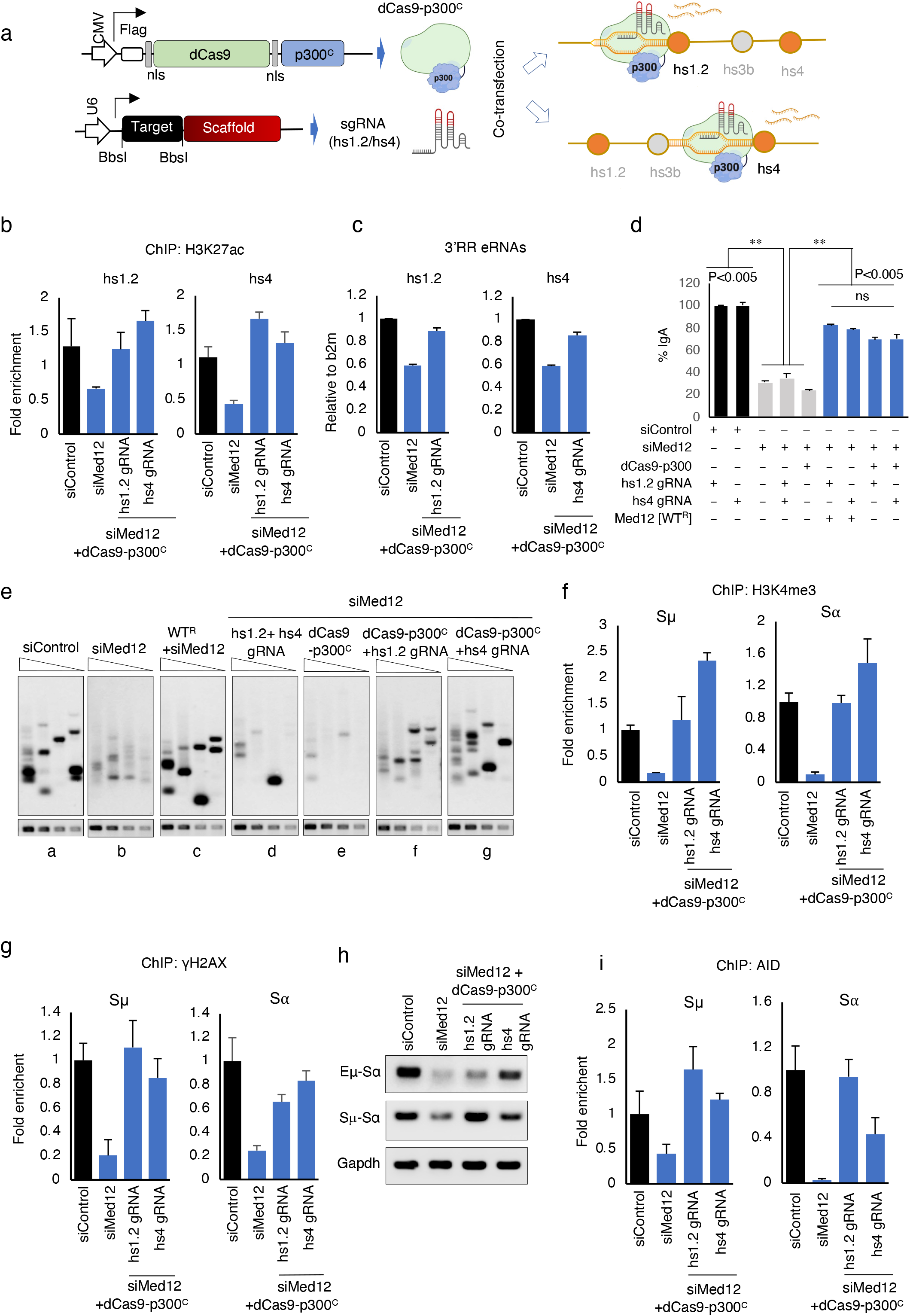
Enhancer targeting CRISPR epigenetic remodeler-activates the 3’RR transcription and fully complemented Med12 deficiency. **a,** The constructs showing the dead Cas9 (dCas9) fused with p300 core HAT domain (dCas9-p300^C^) and expressed from CMV promoter. The small guide RNA (sgRNA) designed for hs1.2 and hs4 were expressed from U6 promoter. The co-transfection of dCas9-p300^C^ and either sgRNAs (hs1.2/hs4) in CH12-F3-2A cells showing site specific enhancer activation at 3’RR. **b,** The H3K27ac ChlP-qPCR showing the enrichment of acetylated H3 histone in dual transfected dCas9+p300C with sgRNAs in Med12KD cells at hs1.2 and hs4 enhancers. The data has been normalized to the input DNA samples followed by maximum values data in each set. The experiments were performed on two independent days and the statistical significance was determined by two-tailed Student’s t-test (P*<0.05). **c**, The RT-qPCR showing the transcripts (hs1.2 and hs4) level in control, siMed12 and dual transfected dCas9-p300^C^ with indicated sgRNAs in Med12^KD^ cells. The data is normalized with ß2m abundance. **d,** The bar plot showing the IgA rescue efficiency in activated and inactivated state of 3’RR enhancers in Med12^KD^ CH12F3-2A cells. The black bar showing the samples treated with control or combination with sgRNAs. The grey bar showing the samples treated with siMed12 or in combination with sgRNAs or dCas9-p300^C^. The left two bar (blue) showing the IgA rescue efficiency in WT-Med12^R^ in Med12^KD^ cells treated with either sgRNAs. The right two bar (blue) showing the recue efficiency in dual transfected dCas9-p300^C^ with sgRNAs (hs1.2 and hs4) in Med12^KD^ cells. The results summarize the means ± s.d. of three independents experiments and the statistical significance was determined by two-tailed Student’s t-test (P*<0.05). **e,** LM-PCR essay estimating the AID induced DNA break in samples treated with dCas9-p300C and/or hs1.2 and hs4 sgRNA alone or in combinations in Med12^KD^ CH12F3-2A cells. The samples have been processed as previously described. **f, g**, The ChIP essay was performed using the indicated antibodies followed by qPCR showing the enrichment at S regions in dual transfected dCas9-p300C with sgRNAs (hs1.2 and hs4) in Med12^KD^ cells. The data has been normalized to the input DNA samples followed by maximum values data in each set. The experiments were performed on two independent days and the statistical significance was determined by two-tailed Student’s t-test (P*<0.05). **h,** The 3C essay showing long range interaction in dual transfected dCas9-p300C with sgRNAs (hs1.2 or hs4) in Med12KD CH12F3-2A cells. Each PCR panel shows the combinations of primer pairs used for DNA amplification. Gapdh PCR of the cross-linked DNA sample served as a loading control. **i,** The ChlP-qPCR showing the AID enrichment in dual transfected dCas9-p300C with sgRNAs (hs1.2 or hs4) in Med12KD CH12F3-2A cells. The values were normalized to the DNA input signals followed by the maximum value in each data set. The experiments were performed on two independent days and the statistical significance was determined by two-tailed Student’s t-test (P*<0.05).

Transfection of dCas9-p300^C^ together with hs-specific sgRNA in Med12^KD^ cells, restored enhancer activity as measured by complete recovery of H3K27ac deposition and a concomitant increase in eRNA transcription (Fig. 5b,c). Interestingly, targeting dCas9-p300^C^ to one of the hypersensitive sites is sufficient to compensate for epigenetic deficits at the other site, possibly due to their palindromic nature and sequence similarity(*58*). As expected, co-transfection of dCas9-p300C and hs-sgRNA rescued the CSR defect in Med12^KD^ cells, nearly as efficiently as by rescue with the WT^R^ construct (Fig. 5d; compare first two blue bars vs last two blue bars). Neither sgRNA (hs1.2 /hs4) or dCas9-p300C transfection alone impacted CSR compensation in Med12^KD^ cells (Fig. 5d; grey bars).

Similarly, co-transfection of dCas9-p300C and hs-sgRNA also restored S region DSB formation in Med12^KD^ cells as detected by our LM-PCR assay (Fig. 5e, compare panel b vs f and g). DSB signals derived from Med12^KD^ cells transfected with hs1.2 and hs4 sgRNA or dCas9-p300C considered as background (Fig. 5e, panels d and e). Strikingly, enforced activation of 3’RR in Med12^KD^ cells also restored S region H3K4me3 and gH2AX marks (Fig.5 f,g). Transfection with dCas9-p300C and hs1.2 or hs4 sgRNA also largely restored long-range interactions between Sμ-Sα/Eμ-Sα in Med12^KD^ cells (Fig. 5h).

As AID is essential to S region DNA DSB and S-S synapse formation, we also examined how enhancer re-activation affects AID recruitment to the IgH locus, a process which has previously been reported to be 3’RR dependent(*14, 17*). In Med12^KD^ cells, AID was indeed reduced at S regions and was restored to control levels upon co-transfection with dCas9-p300C and enhancer-targeting sgRNAs (Fig. 5i). AID has also been reported to target 3’RR and was found to be distributed across the 3’RR super-enhancer region(*59, 60*) Furthermore, AID-induced DNA breaks at 3’RR are responsible for locus suicide recombination (LSR), a process that removes a large sequence region containing Sμ and 3’RR and leads to impaired cell survival. LSR is considered an intrinsic mechanism to regulate B cell homeostasis and for selection of antigen-specific B cells. We observed Med12 depletion also reduces AID recruitment at 3’RR (Extended Data Fig. 4a). However, it remains to be determined whether Med12 and/or 3’RR eRNA also regulate LSR(*59*).

Taken together our findings suggest 3’RR activation by CRISPR/dCas9-p300^C^ compensates for CSR defects following Med12 deficiency, likely through restoring multiple epigenetic processes in the S region, including: H3K4me3 formation, DNA double strand break induction, and S-S synapse formation. In summary, Med12 is an indispensable chromatin remodeling factor for IgH 3’RR activation and associated CSR.

### Mechanism of Med12-dependent IgH 3’RR eRNA regulation

Here we provide evidence that Med12-dependent 3’RR eRNA transcription is critical for CSR regulation. However, several outstanding questions remain, including, how Med12 is recruited to the IgH enhancer and how the enhancer activation functions of Med12 are regulated. To address these questions, we utilized an ER bound enhancer elements (ERE) eRNA transcription system. In this system eRNA transcript expression is strongly and rapidly induced following estrogen exposure(*61*). Recent evidence suggests Jmjd6, a multifunctional enzyme, regulates ERE derived eRNA transcription by recruiting Med12 and the Carm1 (co-activator associated arginine methyltransferase 1) complex(*62*). Jmjd6 may also be necessary for Med12 interaction with Carm1, which in turn methylates Med12 at multiple arginine sites to facilitate chromatin binding.

To test involvement of the Jmjd6/Carm1 pathway in 3’RR regulation through Med12, we first confirmed the presence of Jmjd6/Carm1 at the IgH 3’RR and that this occupancy can be depleted by using siRNA against Jmjd6 (Fig. 6a). We find by ChIP analysis that Jmjd6/Carm1 occupies the 3’RR, and that this occupancy is dependent on Med12 (Fig. 6b). This finding prompted us to examine whether Jmjd6/Carm1 depletion also affects CSR and 3’RR eRNA transcription. Indeed, we find that depletion of Jmjd6/Carm1 reduces CSR as well as eRNA transcription from both hs1.2 and hs4 and is correlated with KD efficiencies (Fig. 6c-d). Furthermore, ChIP analysis revealed that reduced eRNA transcription in Jmjd6^KD^ cells was likely due to loss of H3K27ac mark but not loss of Med12 at the 3’RR, as Jmjd6 depletion specifically reduced H3K27ac (Fig. 6e-f). We confirmed the specificity of siJmjd6 by using an siRNA resistant transcript (WT-jmjd6^R^) that successfully restored the siJmjd6 mediated CSR inhibition (Extended Data Fig. 4d). As expected, expression of WT-Jmjd6^R^ but not the catalytically defective mutant (H187A-Jmjd6^R^) restored CSR in Jmjd6^KD^ cells (Extended Data Fig. 4d), suggesting the demethylase activity of Jmjd6 may remove repressive histone marks to preserve H3K27ac at the 3’RR. To further confirm the observed siJmjd6-mediated CSR impairment is due to loss of H3K27ac and eRNA at the 3’RR, we activated the enhancer by co-transfecting dCas9-p300^C^ and hs4-sgRNA. This activation fully restored the CSR defect in Jmjd6^KD^ cells (Fig. 6g), further emphasizing the importance of 3’RR activation and eRNA production for CSR.

**Fig. 6.**
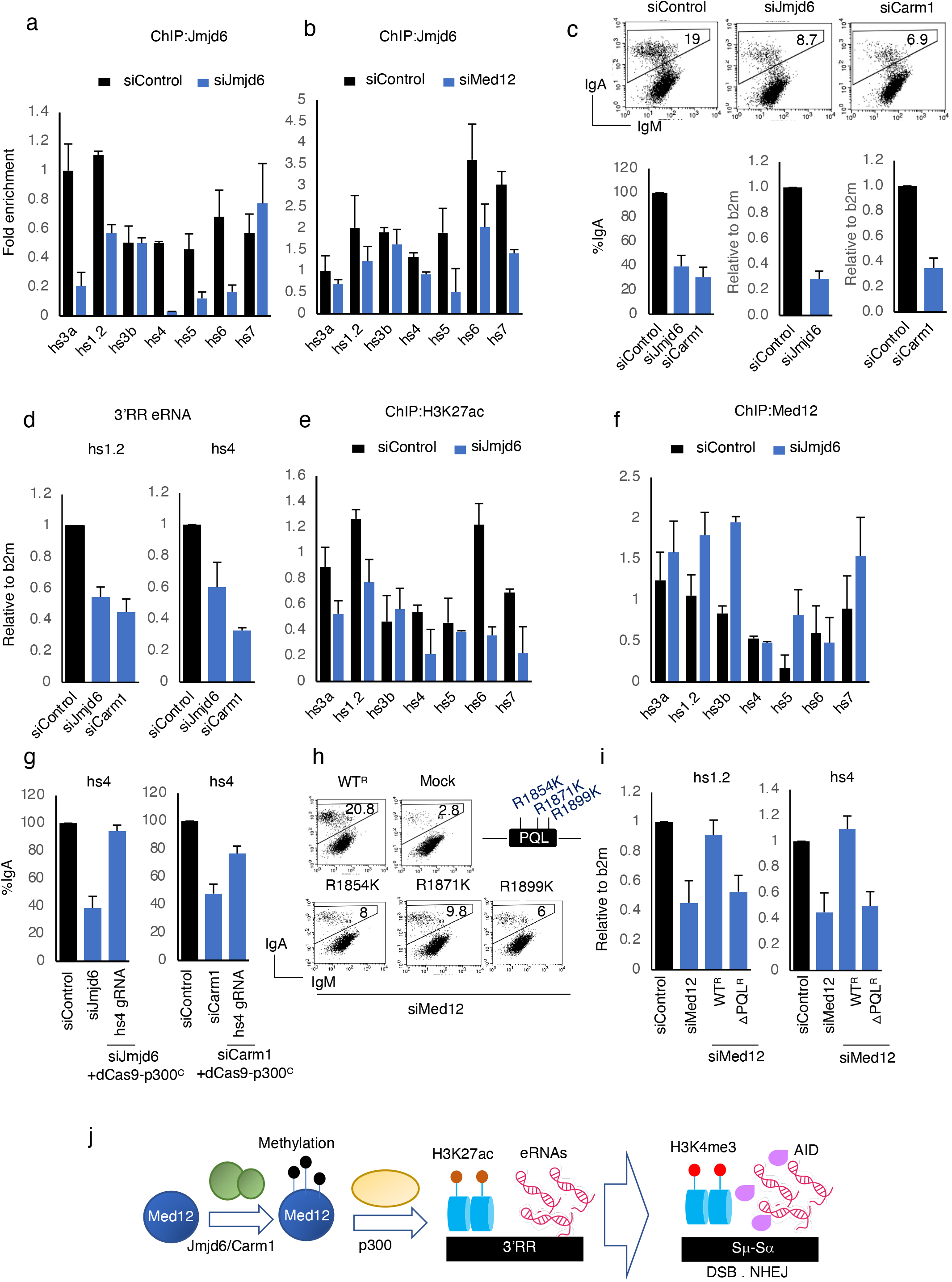
Med12 methylation by Jmjd6/Carm1 initiates enhancer activation. **a,b,** The Jmjd6 ChIP essay was performed followed by qPCR showing the occupancy of Jmjd6 protein at 3’RR in control and either Jmjd6 or Med12 depleted CH12-F3-2A cells. The data has been normalized to the input DNA samples followed by maximum values data in each set. **c,** (Upper panel) The FACS data showing the effect of Jmjd6 and Carm1 knockdown by respective siRNAs in CH12F3-2A cells. The samples were collected after 24 h of CIT (+) stimulation. (Lower panel-left) The RT-qPCR data showing the knockdown efficiency by siJmjd6 and siCarm1 knockdown cells. The data is normalized with b2m abundance. The result summarizes the means ± s.d. of three independents experiments and the statistical significance was determined by two-tailed Student’s t-test. **d,** The bar plot showing the effect of Jmjd6 and Carm1 knockdown on 3’RR hs1.2 and hs4 transcripts. The data is normalized with b2m abundance. **e, f,** The ChIP essay was performed by indicated antibodies followed by qPCR in control and Jmjd6 depleted CH12F3-2A cells.The data has been normalized to the input DNA samples followed by maximum values data in each set. **g,**The bar plot showing the IgA rescue efficiency in dual transfected dCas9+p300^C^ with hs4 sgRNAs in Jmjd6 and/or Carm1 depleted cells. **h,** The FACS plot showing the IgA rescue efficiency in methylation defective Med12 mutants in Med12^KD^ cells. The positions of the mutations at PQL domain are shown at right. **i,** The RT-qPCR analysis showing the rescue of hs1.2 and hs4 transcripts in WT^R^ and PQL (methylation site) deleted Med12 mutant in Med12^KD^ cells. **j,** Schematic representation showing the sequential steps of Med12 workflow in CSR. Med12 is methylated by carm1/Jmjd6 complex (green) at different positions (black boll). Methylation recruits p300 protein to 3’RR which marks histone H3K27 acetylation and activate the enhancers. Activated enhancers transcribed into eRNA which regulates H3K4me3 at S region and recruits DNA break and repair complex for CSR.

Next, to evaluate the importance of Med12 arginine methylation in CSR and 3’RR eRNA regulation, we generated Med12 mutants (R1854K, R1871K and R1899K) that are defective in arginine methylation by Jmjd6/Carm1(*62*). CSR complementation assays showed all three methylation mutants, were defective in CSR, at variable levels of deficiency (Fig. 6h). These CSR defects are comparable to defects previously reported in the Med12 ΔPQL mutant (Fig. 2a-b), the Med12 domain that contains critical and additional arginine methylation sites. Notably, addition of the ΔPQL mutant was also unable to restore 3’RR eRNA transcription from both the hs1.2 and hs4 enhancers in Med12^KD^ cells (Fig. 6i). Taken together, we conclude that the Jmjd6/Carm1 complex is an important post-translational regulator of Med12, which cooperates with p300 to regulate 3’RR activation and eRNA transcription and efficient CSR (Fig. 6j).

### Depletion of 3’RR eRNAs perturbs enhancer function and impairs CSR

Enhancer RNAs are short transcripts that are transcribed unidirectionally or bidirectionally from enhancers(*24*). eRNAs exert a plethora of functions either in *cis* or *trans,* including regulating transcription through recruiting transcription and/or chromatin remodeling factors, regulating long range interaction between enhancer and promoter sequences, and through regulating histone modifications(*63, 64*). At some enhancers, H3K27ac deposition and Mediator complex recruitment are dependent on eRNA production(*64, 65*). From this, we attempted to clarify the role of 3’RR eRNAs in histone modification at the 3’RR and S region, as well as the role of 3’RR eRNAs in S region DSB, S-S synapse formation, DNA repair and CSR.

To specifically deplete 3’RR eRNAs, we designed Locked Nucleic Acid (LNA) modified Antisense Oligos (ASO1 and ASO2) targeting the hs4 enhancer. We chose hs4 due to its basal eRNA expression in CH12F3-2A cells, which is several fold higher than hs1.2 basal eRNA expression (Extended Data Fig. 4b). Transfection of hs4 specific ASOs but not an unrelated control ASO strongly inhibited IgA switching in CH12F3-2A cells (Fig. 7a). Depletion of hs4 specific eRNAs by ASO1 and ASO2 was also confirmed by qRT-PCR (Fig. 7b). Neither μGLT or aGLT transcript expression was affected by ASO1/ASO2 treatment (Fig. 7b). A similar level of CSR impairment by ASOs was also observed in the AIDER expressing CH12-F3-2A cell line, where CSR is induced through activation of an ER fused AID by OHT (tamoxifen) treatment (Extended Data Fig. 4c).

**Fig. 7.**
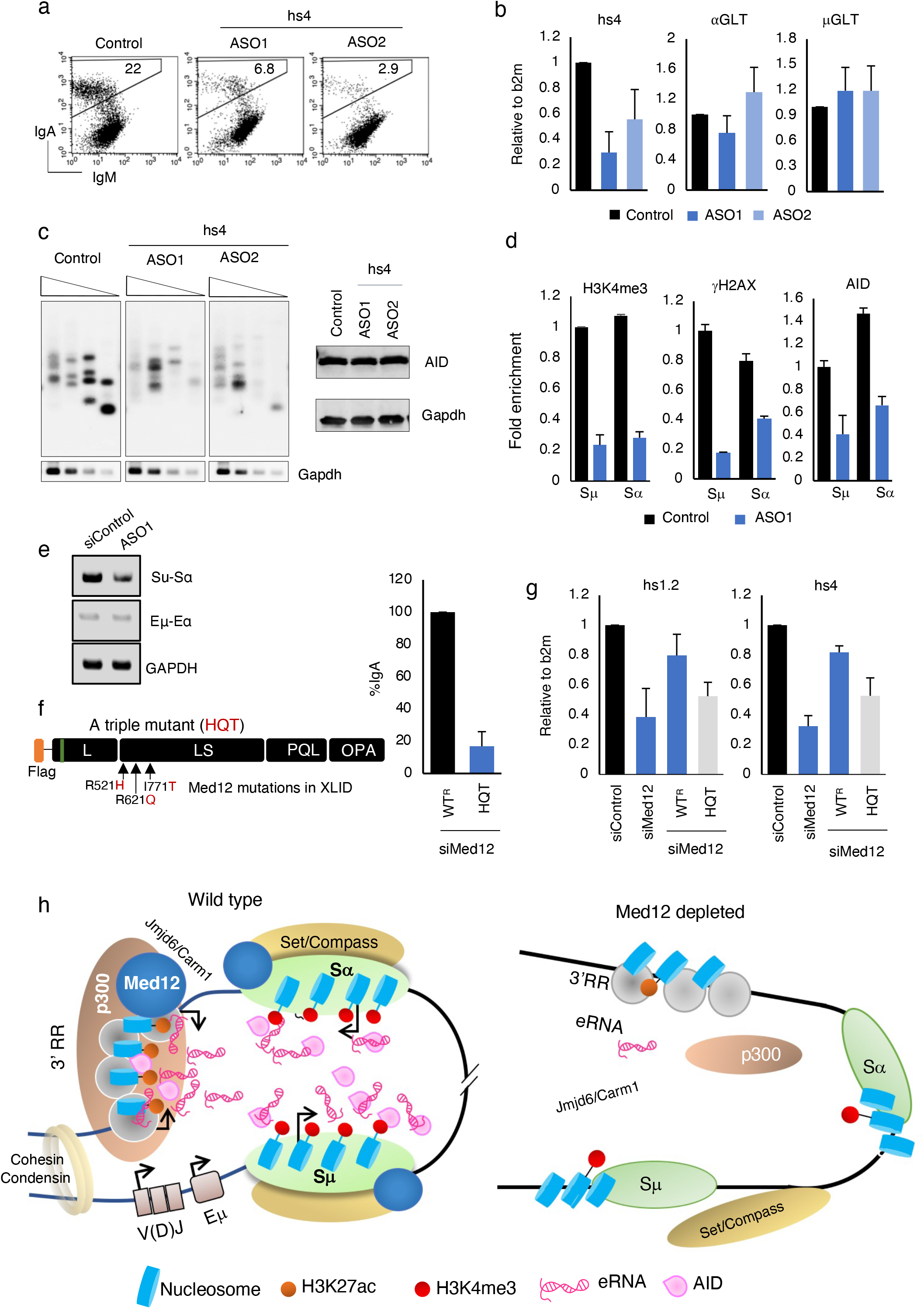
3’RR transcribed eRNA regulates AID induced DNA break and S-S synapsis. **a,** FACS data showing the effect of antisense oligoes (ASOs) designed against hs4 derived enhancer transcripts (ASO1, ASO2), on IgA switching. **b,** RT-qPCR data showing the effect of ASO1 and ASO2 on GLTs and hs4 transcripts relative to ASO control treated cells. The data is normalized with b2m abundance. **c,** (left) LM-PCR essay showing the effect of AID induced DNA breaks in ASO1 and ASO2 treated cells relative to ASO control. Bottom panel shows the semiquantitative PCR analysis of Gapdh of respected samples as an internal control. The triangles indicate a threefold dilution in the DNA amount. (Right) The WB showing the effect of both ASOs on AID protein expression. Gapdh was used as a loading control. **d,** The ChIP essay was performed using the indicated antibodies followed by qPCR showing the enrichment at S regions in control and hs4-ASO1 knockdown in CH12-F3-2A cells. The data has been normalized to the input DNA samples followed by maximum values data in each set. **e,** 3C essay showing the long-range interaction in control and ASO1 knockdown CH12F3-2A cells. Each PCR panel shows the combinations of primer pairs used for DNA amplification. Gapdh PCR of the cross-linked DNA sample served as a loading control. **f,** Schematic representation of Med12 protein showing the mutations found in nonspecific XLID disease. The IgA rescue efficiency has been determined in XLID associated triple mutant (HQT) in Med12^KD^ CH12F3-2A cells. **g,** The RT-qPCR showing the effect of Med12 HQT mutant on 3’RR (hs1.2 and hs4) transcription in Med12^KD^ CH12F3-2A cells. The results summarize the means ± s.d. of three independents experiments and the statistical significance was determined by two-tailed Student’s t-test. **h,** (Right) The proposed model showing the role of Med12 in enhancer activation. Med12 recruits p300 at 3’RR enhancers which in turn catalyzes the H3K27ac histone acetylation. Med12-p300 complex initiates the enhancer activation which transcribed into the eRNA. The produced eRNA act far from the 3’RR and recruits Med12, H3K4me3 methyltransferase and DNA break complex at S regions. (Left) In absence of Med12, p300 doesn’t recruit to 3’RR resulting inactivation of enhancers and perturbed DNA break and S-S synapse.

Next, we examined whether CSR defects are correlated with S region DSB by LM-PCR and ChIP assays. As expected, transfection of ASOs into CH12F3-2A cells dramatically reduced S region DSB signals, as compared to control-ASOs (Fig. 7c). Using the most potent ASO, we evaluated DNA breaks and damage response associated histone marks at the S regions. We observed a severe defect in H3K4me3 deposition and gH2AX formation in ASO1 transfected cells (Fig. 7d), which was consistent with S region DSB impairment (Fig. 7c). AID association with the IgH locus decreased following ASO1 treatment (Fig. 7d), despite AID levels remaining unchanged (Fig. 7c). Notably, in addition to impaired AID-induced S region DNA breaks, long-range Sμ and Sα interactions were also disrupted (but not Eμ-3’RR) following ASO1 transfection (Fig. 7e). It is notable that 3’RR-eRNA depletion following ASO treatment also impaired the two key steps of CSR: S region DSBs and S-S synapse formation-consistent with our observations following Med12 depletion. Taken together, our data suggest Med12 functions in CSR are dependent on eRNA expression from the 3’RR.

Interestingly, we find the Med12 mutant with multiple XLID mutations (R521H/R621Q/I771T), which we named “HQT”, shows significant CSR impairment and 3’RR eRNA transcription defects (Fig 7f-g). Such a function has never been reported for any XLID mutations; we also find loss of the Med12 PQL domain impairs CSR and 3’RR eRNA transcription (Fig. 2b, Fig. 6i). Our findings that the LS and PQL domains of Med12 regulate CSR implies that both AID-driven DNA DSBs and S-S synapse formation require 3’RR activation-and more specifically eRNA transcription. From this, we propose IgH 3’RR derived eRNAs drive epigenomic regulation at the IgH enhancer and S-region, which stabilizes S-S proximity and promotes AID-dependent DNA double strand breaks and CSR (Fig. 7h).

## DISCUSSION

### Med12 and AID have functional homology for DNA double strand breaks and S-S synapse formation during CSR

AID is an indispensable enzyme for generating the DNA double strand breaks and S region synapses that are required for oriented deletional recombination during *CSR*(*10, 11, 66–68*). How AID regulates the mechanistically distinct processes of S region DNA double strand breaks and S-S synapse formation remains poorly understood(*69*). While numerous AID-interacting and non-interacting CSR regulatory factors have been identified, these factors appear to play exclusive roles in either DNA break, repair, or synapse formation(*4, 70–74*). Here, we show Med12 is a novel CSR cofactor that not only promotes AID-induced DNA double strand breaks and synapse formation but also supports DNA repair via NHEJ (Fig. 1d-f, Extended Data Fig. 1e). We find that Med12 shares with AID many of the key functional aspects that are essential for efficient CSR and eventual Ig gene diversity. As CSR is highly sensitive to Med12 protein levels, we consider Med12 levels to be a key rate-limiting factor for DNA double strand breaks and S region synapse formation. Our analysis of multiple Med12 mutations (Extended Data Table 1) shows the LS and PQL domains are critical for efficient CSR (Fig. 2e) and likely have distinct, yet overlapping functions. Loss of the LS, but not PQL domain, leads to severe impairment of DNA double strand breaks and DDR signals such as gH2AX formation in the S-region (Fig. 1d,e, Fig. 2c). However, deletion of either domain leads to impaired S-S synapse formation and loss of AID occupancy at the IgH locus (Fig. 2d, Fig 5i). Our cross-complementation studies using Med12 mutants also suggests that while the LS domain is crucial for AID-induced DNA double strand breaks both the LS and PQL domains are necessary for S-S and Eμ-3’RR interaction (Fig. 2d, Fig. 1f).

### Med12-dependent IgH 3’RR super-enhancer activation is central to CSR

Consistent with Med12’s function in IgH locus DNA double strand breaks and long-range chromatin looping, we find Med12 enrichment at the Eμ, Sμ, Sα, and 3’RR super-enhancer regions(*75*) (Extended Data Fig. 5ab,c). However, Mediator is a 30-subunit megadalton complex with four sub-modules, including the Med12-containing kinase module that reversibly interacts with core Mediator for transcriptional regulation(*24, 26*). Interestingly, at the IgH locus Med12 functions in CSR appear independent of its kinase function, as CSR was refractory to CDK8 depletion or catalytic inhibition of the CDK8 kinase. These findings are also consistent with competent CSR observed in Med12 mutants that are defective in CDK8 activation (Fig. 3f,g). Similarly, depletion of Med13/Med13L, the essential bridging component between core Mediator and the kinase module, does not affect CSR (Fig. 3h). While novel associations may still exist, these kinase-independent functions of Med12 are reminiscent of Med12 functions in HPSC homeostasis(*27*), where Med12 helps regulate cell lineage specific super-enhancers in a kinase-independent manner.

H3K27ac and H3K4me1 on enhancer chromatin distinguish transcriptionally active enhancers from inactive and paused enhancers(*76–78*). Priming of enhancer activation occurs through the binding of pioneering TFs and subsequent loading of other TFs, coactivators, and RNAPII, to generate eRNA from super-enhances(*79*). Consistent with previous reports, our work suggests Med12 maintains IgH 3’RR super-enhancers activation through p300, as Med12 depletion results in p300 recruitment defects, impaired H3K27ac deposition and 3’RR eRNA transcription (Fig. 4d,c,i,j). Here, we used multiple lines of evidence to demonstrate the importance of IgH enhancer activation for CSR. These lines of evidence included: (1) a Med12 deficiency complementation for CSR by CRISPR/dCas9 enhancer activation system that restored H3K27ac signature and eRNA synthesis at 3’RR (Fig. 5b,c), (2) recapitulating 3’RR inactivation and CSR defects observed in Med12 deficiency by p300 depletion or inhibition (Fig. 4c,j,k), (3) degradation of 3’RR eRNAs by ASO targeting shows strong CSR impairment, as well as impaired S region DNA double strand breaks, reduced long-range interactions and reduced AID recruitment to the IgH locus (Fig. 7c,e,d). We obtained similar results when eRNA processing was interrupted by depleting the catalytic subunit of the integrator complex (IntS11,13) (Extended Data Fig. 4e), which regulates the length and production of eRNAs(*80, 81*). Consistent with these findings, we also find enrichment of IntS11 and IntS13 at the 3’RR enhancer after stimulation with CIT for CSR (Extended Data Fig. 4f).

The requirement of the 3’RR super enhancer in CSR has been well documented by numerous elegant studies in mice models and *ex vivo* system(*13, 17, 82*). However, there remains minimal information regarding the mechanisms of 3’RR activation and how this activation affects AID-induced CSR events. Recently, Zymnd8 that recognizes various histone modifications was shown to reduce RNAPII loading at the IgH super-enhancer and reduce enhancer transcription(*83*). Conversely, RNAPII elongation factor Spt5 may regulate 3’RR transcription by regulating RNAPII pause release(*84*). Spt5 deficiency does not disrupt Med12 or H3K27ac occupancy at the 3’RR, and a CRISPR/dCas9-VPR transcriptional activation system can rescue enhancer transcription defects but not CSR. This is likely due to Spt5 being required for AID expression and recruitment(*84, 85*). However, we find that complementation of Med12 deficiency strictly requires an enhancer activating CRISPR/dCas9-p300^C^ system, which restores H3K27ac marks at the 3’RR and restores eRNA transcription and CSR (Fig. 5d). In conclusion, our results are consistent with Spt5 and Zymnd8 functioning downstream of enhancer activation by Med12.

### Med12 and 3’RR eRNA dependent epigenomic and conformational regulation

Promoters of actively transcribed gene loci are generally enriched with H3K4me3, a signature mark of accessible chromatin(*86, 87*). However, at the IgH locus actively transcribed I promoters and downstream S recombination regions are highly enriched with *H3K4me3*(*49, 50, 52, 88–91*). Downregulation of IgH locus-specific H3K4me3 is associated with various CSR-associated defects, including impaired AID-induced S region DNA double strand breaks and impaired Ig isotype-specific transcription in primary B cells(*49, 50, 92*). Our work here is consistent with other studies suggesting that the combinatorial histone code, including H3K4me3, facilitates DNA double strand break complex formation at the S region(*49, 50, 52, 88, 93*). One of our striking findings is that Med12 deficiency causes strong downmodulation of H3K4me3 as well as AID-induced S region DNA double strand breaks. As Med12 is in a complex with H3K4me3 writing machinery, loss of Med12 in S regions may perturb H3K4me3-complex activity, leading to impaired AID-induced DNA double strand breaks. Within this process, we suspect 3’RR activation plays a critical role, as p300 depletion (or 3’RR inactivation) leads H3K4me3 impaired deposition and DNA double strand break formation at S regions (Fig. 4h,f). This hypothesis is consistent with our finding that 3’RR activation by CRISPR/dCas9-p300 system can fully restore defects caused by Med12 depletion.

Most importantly, depletion of 3’RR eRNAs, the activation markers of the IgH superenhancer, phenocopied the DNA double strand break and S-S synapse formation defects observed in Med12, p300, and AID deficiency (Fig. 7c,d,e). This result suggests a direct and unique role of 3’RR eRNAs in CSR regulation. We have shown that 3’RR eRNAs can bind Med12, Wdr5 and Rad21 (Extended Data Fig. 5d), and their depletion impairs recruitment of Med12, AID, and H3K4me3 occupancy at S region, leading to DNA double strand break and S-S synapse formation defects (Extended Data Fig. 5c, Fig. 7d), Notably, 3’RR eRNA depletion did not affect Med12 or H3K27ac occupancy at the 3’RR, suggesting these eRNAs predominantly contribute to CSR in a trans manner, possibly functioning in the S regions. Consistent with this hypothesis, S region DNA double strand breaks and long-range Sμ-Sα but not Eμ-3’RR interactions are affected by 3’RR eRNA depletion. However, Med12 depletion leads to disruption of both chromatin loops, Sμ-Sα and Eμ-3’RR interactions (Fig. 1f), suggesting Med12 is an essential constituent of both Sμ-Sα and Eμ-3’RR chromatin conformations. While basal levels of Med12 and 3’RR eRNAs are likely sufficient to confer Eμ-3’RR chromatin architecture, CSR induction may stabilize it further through Med12 enrichment and increased eRNA transcription. eRNA transcripts also may help Med12, AID, and other essential components localize to recombining S regions for S-S synapse formation.

### Involvement of 3’RR eRNAs in CSR-specific condensate formation

Recent studies have shown super-enhancers exist as phase-separated condensates, defined as a 3D reservoir holding large-scale regulatory factors and RNAs(*94–96*). Associated eRNAs have been implicated in stimulus-dependent(*81, 97*) (e.g. E2/EGF) dynamic phase separation during enhancer-promoter looping where they interact with many factors present in super-enhancers, including Med1, Cohesins, YY1, p300/CBP, and BRD4 (*64, 98–100*). Depletion of eRNAs from associated enhancers affected either the recruitment or function of the chromatin conformation regulatory factor, leading to disruption of enhancer-promoter chromatin looping and/or enhanceosome formation(*94, 95, 99, 101*). Our work also shows that depletion of 3’RR eRNA significantly affects recruitment of Med12, AID and possibly still unknown components to the S region. PONDR analysis (Extended Data Fig. 6a) suggests Med12 has intrinsically disordered region (IDR) as observed in Med1 and Brd4, both of which promote enhancer condensate formation. We also confirm that Med12, like Med1 and Brd4, can be precipitated by IDR specific chemical b-isox(*102*) (Extended Data Fig. 6c). Additionally, eRNA transcription and CSR are both sensitive to 1,6-HD treatment, a known method for perturbing eRNA-dependent enhancer ribonucleoprotein complex (eRNP) formation and chromatin architecture (Extended Data Fig. 6b)(*94, 103*).

From this, we suspect 3’ RR-derived eRNAs can act as a matrix or recruiting medium so DNA break-recombination complexes can assemble locally. While some of these defects have been previously observed in a 3’RR mouse model, the underlying mechanisms remain unclear. Our work here provides a plausible explanation for how enhancer activation and transcription can coordinate two seemingly dissimilar processes simultaneously, AID-induced DNA double strand breaks and S-S synapse formation.

### Implications for Med12 and dysregulated CSR in X-linked intellectual disability syndrome

Here, we have shown that mutations located near the N-terminus of Med12 and those affecting Med12 kinase module(*48*) functions do not impact CSR. Conversely, several LS and PQL domain-associated Med12 mutations show significant impairments in CSR (Fig. 3g, Extended Data Table 1). Both atypical and FG-type XLID mutations showed a dramatic impairment of CSR with impaired 3’RR activation/transcription (Fig. 7f,g). It remains unknown whether patients with these XLID mutations(*104, 105*) display mild to moderate B cell abnormalities or have Ig deficiency. Our work here suggests the LS and PQL domains of Med12 both coordinate regulation of eRNAs (Fig. 7g, Fig. 6i), the markers of active 3’RR super-enhancers. Although, Med12 mutations have been linked to various cancer types and congenital disorders with intellectual disability, the underlying functional defects and mechanisms of these disorders remains poorly understood(*32*) (Extended Data Table 1). Defined and coordinated regulation of super-enhancers is essential for cell lineage and cell identity(*27, 106, 107*). From this, it is conceivable that Med12 defects lead to dysregulation of normal/ or cell lineage-specific super-enhancer functions, which ultimately give rise to neurologic disorders and/or tumors(*106, 108, 109*). Furthermore, many neurodevelopmental and/or intellectually disability genes are linked to DNA damage repair and the NHEJ pathway(*110–112*). As NHEJ pathway is essential for both V(D)J and CSR, DNA repair defects are often a common cause of both neuronal and B cell defects. We show Med12 deficiency may impair NHEJ; however, additional work will be needed to determine the mechanism.

### Methylated Med12 initiates enhancer activation

It remains unclear how Med12 coordinates these molecular activities. It is possible that Med12’s functions in CSR are regulated by interacting with other noncoding RNAs, specifically eRNAs and lncRNAs (unpublished) and/or through arginine C-terminal methylation to regulate protein-protein or protein-RNA interactions. Carm1 methylates multiple arginine residues at the C-terminus of the Med12 PQL domain. Carm1-dependent Med12 methylation sensitizes breast cancer cells to chemotherapy and mutations at these methylation sites impair Med12 binding at the p21 locus(*36*). Recent work has shown Jmjd6 regulates Med12 methylation through Carm1, which promotes Med12 binding to ERa bound active enhancers and regulates Med12 function(*62*). Our results partially support this model, as we find depletion of the methylation/demethylation complex (Jmjd6/Carm1) also reduces eRNA transcription and CSR (Fig. 6d,c). Interestingly, we find Med12 is required for recruiting the Carm1/Jmjd6 complex to the IgH 3’RR locus, while Carm1/Jmjd6 are dispensable for Med12 recruitment. Furthermore, depletion of the demethylase Jmjd6 results in failures of enhancer activation (Fig. 6e). From this, we suspect the methylation status of Med12 is crucial for enhancer activation (possibly through methylation-dependent recruitment of other coactivators/histone modifiers such as p300) to the IgH 3’RR locus, while methylation status does not affect Med12 chromatin binding. This hypothesis is strongly supported by evidence that Carm1 methylates Med12 at R1899, which is then recognized by coactivator TDRD3, leading to further interactions with activating-ncRNAs for estrogen regulated gene transcription(*35*). Crucially, we find R1899 mutant Med12 also has diminished CSR activity (Fig. 2e). We did not find any significant difference in eRNA transcription in individual Med12 methylation defective mutants (data not shown). However, deletion of the entire PQL domain of Med12 leads to defective eRNA regulation (Fig. 6i), suggesting combinatorial methylation is likely required for enhancer activation, a conclusion that is consistent with a previously published report(*62*).

## Acknowledgements

This work was supported by funding from the Japan Society for the Promotion of Science; Grant-in-Aid for Scientific Research (S) 15H05784 to T.H. and Grant-in-Aid for Scientific Research (C) 21K06015 to N.A.B. We would like to thank Dr. M.D. Shair and Dr. D.J. Taatjes for providing the Med12 kinase (CDK8 and CDK19) inhibitor. F.H would also like to specially thanks to his wife Samina for her moral support and his mentor T.Honjo for arranging and providing the funds. We also would like to thank Ms. M. Nakata for technical assistance.

## Data availability

This study includes no data deposited in external repositories.

## Author contributions

F.H. and N.B. designed the experiments and wrote the manuscript. F.H performed the experiments. T.H and N.B. conceptualized and supervised the project. F.H, N.B and T.H reviewed the manuscript.

