## Supplementary for "XLID Syndrome Gene Med12 Promotes Ig Isotype Switching through Chromatin Modification and Enhancer RNA regulation"

### Material and Methods

#### Cell culture, siRNA Transfection and CSR

For CSR, a derivative of mouse B-cell lymphoma line (CH12F3-2A) expressing Bcl2 was used throughout the study(*1*) unless stated. Cells were cultured and maintained in RPMI supplemented with glutamine, NCTC, FBS (10%), β-mercaptoethanol, and penicillin/streptomycin. For knockdown experiments, the Silencer/Stealth siRNA or control (low GC) oligonucleotides were purchase from Thermo Fisher Scientific, MA, USA and the antisense oligoes (ASO) (control or target) were purchased from QIAGEN, were introduced into the cells using the Nucleofector 96-well electroporation system (Lonza, Switzerland) as described by manufacturer instructions. After the transfection, the cells were cultured for 24 h and then were stimulated by CIT cocktail(*1*) (anti-CD40L, IL4, and TGFβ) or OHT (1 mM) for AIDER cells to induce IgM to IgA isotype switching, and cultured for another 24 h-48 h before collection.The surface expression of IgM and IgA was examined by staining the cells with FITC-conjugated anti-mouse IgM (eBioscience) and PE-conjugated anti-mouse IgA (eBioscience). Propidium iodide (PI) staining was included to stain the dead cells. The FACS analysis was performed with a BD FACS Calibur instrument, and the data were analyzed by CellQuest software (BD Biosciences). All of the sequences of the siRNA oligos are shown in SI appendix.

#### Plasmids

The mouse Med12 (NM_021521) cDNA was obtained by RT-PCR of the RNA isolated from CH12F3-2A cells and cloned into the *AsiS*I/*Mlu*I (Takara) sites of the pCMV6-Entrymammalian expression vector (Origene- PS100001).The 3xFlag epitope was fused at the N-terminal of Med12 during the RT-PCR to express 3xFlag-Med12 wild-type construct. To generate siMed12-resistant Med12 transcripts ($WT^R$-Med12), the siMed12 targeting sequence (CCAUCUACUGUAACGUGGA) was modified to (CGATATATTGCAATGTAGA) without altering the encoded amino acids. $WT^R$-Med12 construct was further used for the deletion or point mutagenesis using the Q5 Site-Directed Mutagenesis Kit (NEB-E0554) according to the manufacturer's instructions.

The mouse Jmjd6 (NM_001363363) cDNA was obtained as described above and cloned into the *Hind*III/*BamH*I (Takara) sites of the pCMV10 vector. The 3xFlag epitope was tagged at N-terminal. To generate siJmjd6-resistant Jmjd6 transcripts (Jmjd6-$WT^R$), the siJmjd6 targeting sequence is modified by multiple mutagenesis without altering the amino acid code. The primers sequence has been provided in the SI appendix.

**Western Blotting (WB)**

For immunoblotting, total cell lysate from were prepared by lysing $2\times10^6$ cells by the RiboCluster Profiler Kit (MBL-RN1001) supplemented with EDTA-free protease inhibitor cocktail (Roche) and RNAase Inhibitor (Sigma). After lysis for 5 min at 4ºC, cells were centrifuged (135,000g) for 5 min and supernatant were collected in separate tube. The obtained clear lysate was mixed with 2xSDS sample buffer in 1:1 ratio and heated for 85ºC for 15 min. The denatured samples were separated by electrophoresis in 4–20% SDS–PAGE (Bio-Rad) and subjected to standard western blot analysis.

**Immunoprecipitation (IP)**

For each IP, Protein G Dynabeads (Invitrogen-1003D) were pre-incubated with antibody of interest (5µg) with rotation at RT for 1 h. Beads were collected and washed 2x with wash buffer provided in a RiboCluster Profiler Kit (MBL-RN1001) before overnight incubation with lysate. After the incubation, beads were washed 3x and add equal ratio of 2xSDS and wash buffer before gel electrophoresis.

**ChIP-qPCR**

The chromatin immunoprecipitation (ChIP) essay was performed using the ChIP-IT Express Kit (Active Motif-53008) according to the manufacturer's instructions. In brief, $5\times10^6$ cells were fixed in the presence of 1% formaldehyde for 5 min at room temperature. The reaction was stopped by the addition of 0.125M glycine. A soluble chromatin fraction containing fragmented DNA of 500–2,000 bp was obtained after cell lysis and sonication. ChIP was performed by incubating the cleared lysate with 3 µg of antibodies. The immunoprecipitated DNA was analyzed by real-time PCR, and the data were normalized first to the amount of input and then to the maximum value in each dataset, as described previously(*2*).The ChIP antibodies and primers used are listed in SI appendix.

**RNA Immunoprecipitation (RIP)**

For RIP assay, $1\times10^7$ cells were collected, washed with cold PBS and lysed by RiboCluster Profiler Kit (MBL-RN1001) supplemented with DTT, RNAase Inhibitor (Sigma) and EDTA-free protease inhibitor cocktail (Roche) as described by the manufacturer. The subsequential RIP was performed using the Magna RIP kit (17-700 Merck Millipore) according to the manufacturer's instruction with slight modifications. After overnight RIP, the TRIzol™ (ambion) was directly added to the washed magnetic beads containing RNAs and the total RNA was extracted using the standard RNA isolation protocol. The purity and concentration of RNA were analyzed by nanodrop or Agilent bioanalyzer 2100.

**RNA Isolation, Reverse Transcription and RT-qPCR**

RNA was isolated from transfected cells using the TRIzol™ (ambion) followed by *DNase* I treatment (SigmaAldrich-AMPD1). 1μg of purified RNA was used as template for complementary DNA synthesis using SuperScript IV (Invitrogen-18090050) with random primers Invitrogen-N8080127). Real-time PCR (Applied biosystem-7900HT Fast Real time PCR System) was performed using SYBR Green Master Mix (Applied Biosystems) and mRNA-specific primers listed in SI appendix. The $2^{-\Delta\Delta Ct}$ method was used to quantify the data using either Gapdh or β2m housekeeping transcript for normalization.

**LM-PCR and 3C essay**

For double stranded DNA break estimation, CIT (+) stimulated $1\times10^6$ cells/sample were collected as described above and fixed in low-melt agarose plugs and processed for linker ligation as described previously(*3*). Briefly, linker ligation reaction was performed overnight at 16°C, and to inactivate the reaction, samples were heated at 70°C for 10 min. 3-fold serial dilutions of linker ligated DNA were amplified by KOD-FX-Neo polymerase (Toyobo) using an Sμ-specific primer (forward) and a linker-specific primer (reverse). To confirm the amplification of Sμ-specific DSBs, Southern blot analysis of the PCR products was performed using a DIG-labeled Sμ probe. Similarly, PCR of the Gapdh locus from each sample served as a normalization control.

The 3C essay was performed as described previously(*4, 5*). Briefly, $7\times10^6$ cells were collected, washed and cross-linked for 5 min at room temperature with 1% formaldehyde. After crosslinking the cells were subjected to nuclear lysis. The cross-linked chromatin was digested overnight with *Hind* III and later ligated with T4 polymerase DNA ligase (Takara).The ligated chromatin was treated with proteinase K and reverse cross-linked, and the DNA was purified by phenol/chloroform extraction protocol. PCRs were performed using the primers described in SI appendix.

#### *IgH/c-Myc* Translocation and NHEJ essay

For translocation essay, genomic DNA has been isolated from 48 h CIT (+) stimulated samples and were PCR amplified at *IgH/c-Myc* translocation junctions using Expand long template PCR system as described previously(*6*). The PCR products were separated by electrophoresis on ethidium bromide-containing 1% agarose gels and subjected to Southern blotting with a Myc-specific probe.

The NHEJ essay was performed as described previously(*7*). In brief, The I-*Sce*I expressing plasmid (pCBASce) alone or with siMed12 (designed against human Med12) or control were co-transfected into NHEJ-reporter cell lines using Lipofectamine 2000 (Invitrogen). Transfected cells were harvested after 48 h before FACS analysis. The NHEJ-reporter cell line H1299dA3-1 were a kind gift from Dr. T. Kohno at the National Cancer Center Research Institute, Tokyo. The primers and siRNAs sequence were shown in SI appendix.

#### Inhibitors

Cortistatin A (CA) was a kind gift from Dr Matthew D. Shair (Harvard University) and Dylan J. Taatjes (University of Colorado Boulder). The dissolved CA in DMSO was provided as a stock concentration of 1mM and was stored at -80ºC for further use. The final concentrations 100 nM and 250 nM were used in the experiments. The DMSO concentrations 0.0001% and 0.00025% (without CA) was used as a vehicle control.

The P300 HAT inhibitor, C646 (Sigma-328968-36-1) was dissolved in DMSO and the 5µM concentration was used for related experiments. The lists of antibodies, primers, and stealth siRNAs used in this study are shown in SI appendix.

#### CRISPR and sgRNAs Cloning

**CRISPRa-**To activate the 3'RR IgH enhancer we exploited the CRISPRa technology(*8*). The sgRNAs (hs1.2 and hs4) sequence was obtained from previously published report(*9*). Briefly, oligonucleotides were annealed in the following reaction: 10 µM guide sequence oligo, 10 µM reverse complement oligo, 50 mM NaCl in Tris-EDTA buffer with the cycling parameters of

95 °C for 2 min and then ramp down to 25 °C at 5 °C/min. The annealed oligos were cloned into the sgRNA vector (pLH-spsgRNA2-Addgene#64114) using a Golden Gate Assembly strategy including: 100 ng of circular sgRNA vector plasmid, 0.2 μM annealed oligos, 20 U of *Bbs*I restriction enzyme, 750 U of T7 DNA ligase (Enzymatics-L6020L) and 1x T7 ligase buffer with the cycling parameters of 37 °C for 5 min followed by 60 °C for 5 min. Insertion of sgRNA was validated by Sanger sequencing (3130xl Genetic Analyzer). The pcDNA-dCas9-p300 Core was purchased from Addgene#61357. Briefly, 500 ng of dCas9 expression vector and 500 ng of individual gRNA expression vectors were transfected in $Med12^{KD}$ cells and incubate the cells for 48 h. After the incubation, the cells were CIT (+) for another 48 h before collecting the cells used for subsequent analysis.

**CRISPRi-**To knockdown the endogenous Integrator complex components (IntS11, IntS13), sgRNA sequence (2 sets for each) were designed from (http://crispor.tefor.net/). The obtained sgRNA oligonucleotide were annealed as described above and cloned in pSPgRNA purchased from Addgene#47108 using the *Bbs*I cloning site. The vector dCas9-KRAB-MeCP2 were purchased from Addgene#11082. Briefly, 500 ng of dCas9 expression vector and 500 ng of equimolar pooled sets of gRNAs were transfected and incubate the cells for 24 h. After the incubation, the cells were CIT (+) for another 24 h before collecting the cells used for subsequent analysis.

**Biotinylated Isoxazole-Mediated Precipitation**

Biotinylated Isoxazole (b-Isox) mediated protein precipitation was performed as described by previously published reports(*10*). The b-Isox was purchased (Sigma-T51161), dissolved in DMSO and added to the cell lysates at 100 or 250 μM final concentrations. The reaction solutions were incubated at 4°C for 1 h and then centrifuged at 14K rpm for 15 min. The pellet was washed twice with the wash buffer and resuspended in 2xSDS sample loading buffer. After SDS-PAGE, protein was detected by western blotting using indicated antibodies. The antibodies used in this study are listed in SI appendix.

**Statistical Analysis**

Error bars represent standard deviation (SD) from either independent experiments or independent samples. All statistical analyses were performed using GraphPad Prism, and the information about statistical methods is specified in figure legends. The numbers of independent experiments or biological replicate samples and P values (n.s. not significant, *P < 0.05) are provided in individual figures. P < 0.05 was considered statistically significant.

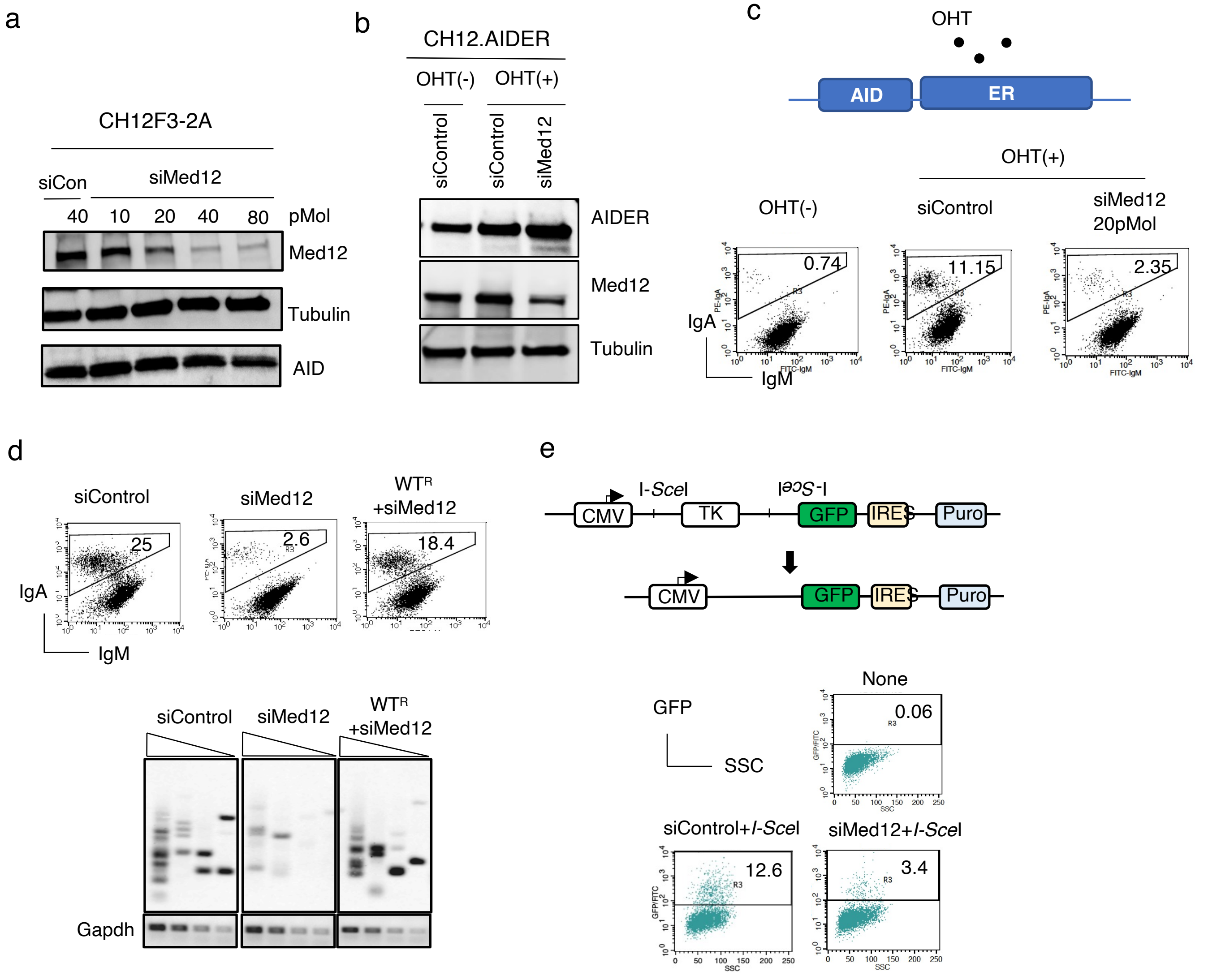


**Extended Data Fig. 1**

**Med12 depletion leads to impaired critical steps of CSR without effecting AID expression.a,** The WB showing the knockdown efficiency of different concentrations of siMed12 on endogenous Med12 transcripts. The CH12F3-2A cells were transfected with increase doses of siMed12 for 24h and CIT(+) for another 24h before harvest the cells for analysis. The blot was probed against the antibody indicated. a-Tubulin was used as internal control.**b,c** CH12F3-2A derived AID overexpressing line where AID is fused with ER (AID-ER) is transfected with siMed12 (20pMol) and later stimulated for 24h with tamoxifen OHT (+) to determine the effect of $Med12^{KD}$ on AID expression and corresponding IgA. The blot was probed against the indicated antibodies. **d,** The FACS analysis estimating IgA switching and corresponding LM-PCR estimating AID induced DNA break in control or siMed12 depleted and exogenously expressed $WT^{R}$ Med12 construct in Med12 depleted CH12F3-2A cells. The cells were harvest after 24h of CIT(+) stimulation and processed as described above. Gapdh was used as internal control.**e,** The schematic showing I-*SceI* induced DNA DSB repair by NHEJ by the reporter construct. The DSB generated by I-*SceI* enzyme removes the intervening thymidine kinase gene and successful joining of the broken end by NHEJ was estimated by GFP expression. The cells transfected in combination with I-*SceI* plasmid either with siControl or siMed12. The GFP positive cells were analyzed by FACS analysis after 24h of transfection.

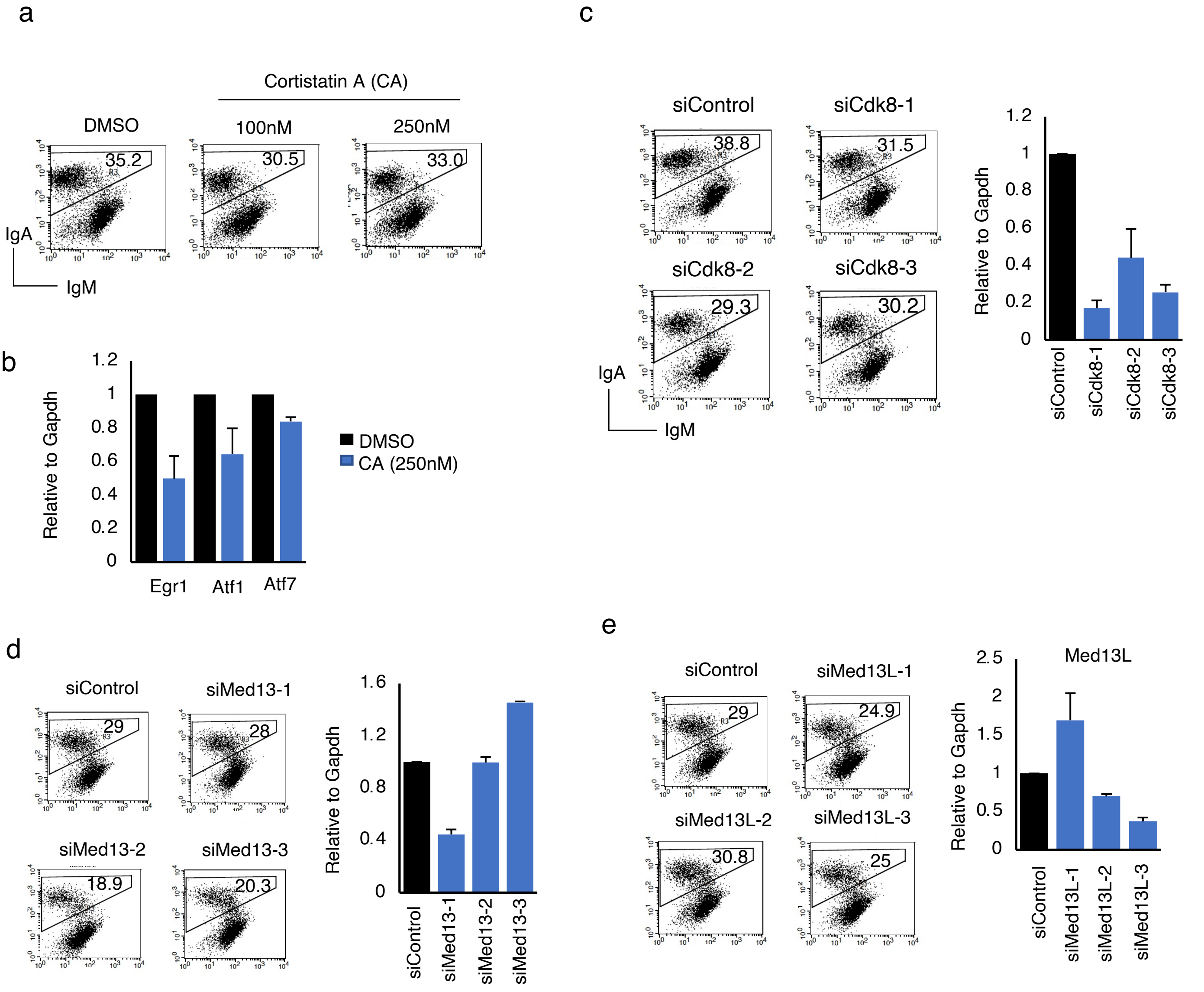


**Extended Data Fig. 2**

**The Kinase module components other than Med12 are not required for CSR.a,** The FACS analysis showing the effect of Cdk8 kinase inhibitor, CortistatinA (CA), on CSR. The stock concentrations of CA were made in DMSO and DMSO without CA used as negative control. The CH12F3-2A cells were stimulated for 48h before analysis.**b,** The RT-qPCR data showing the effect of CA (250nM) on transcripts known to inhibited by CA. DMSO without CA used as negative control. The data is normalized with gapdh abundance.**c,d,e,** Three different sets of siRNAs transfected against the Cdk8, Med13 and Med13L and checked their effect on CSR after 48h of CIT(+) stimulation in CH12F3-2A cells. The corresponding KD efficiency by each siRNAa was measured by RT-qPCR analysis. The data is normalized with Gapdh abundance. The result summarizes the means ± s.d. of three independents experiments and the statistical significance was determined by two-tailed Student’s t-test (P>0.05), ns indicates in significant difference.

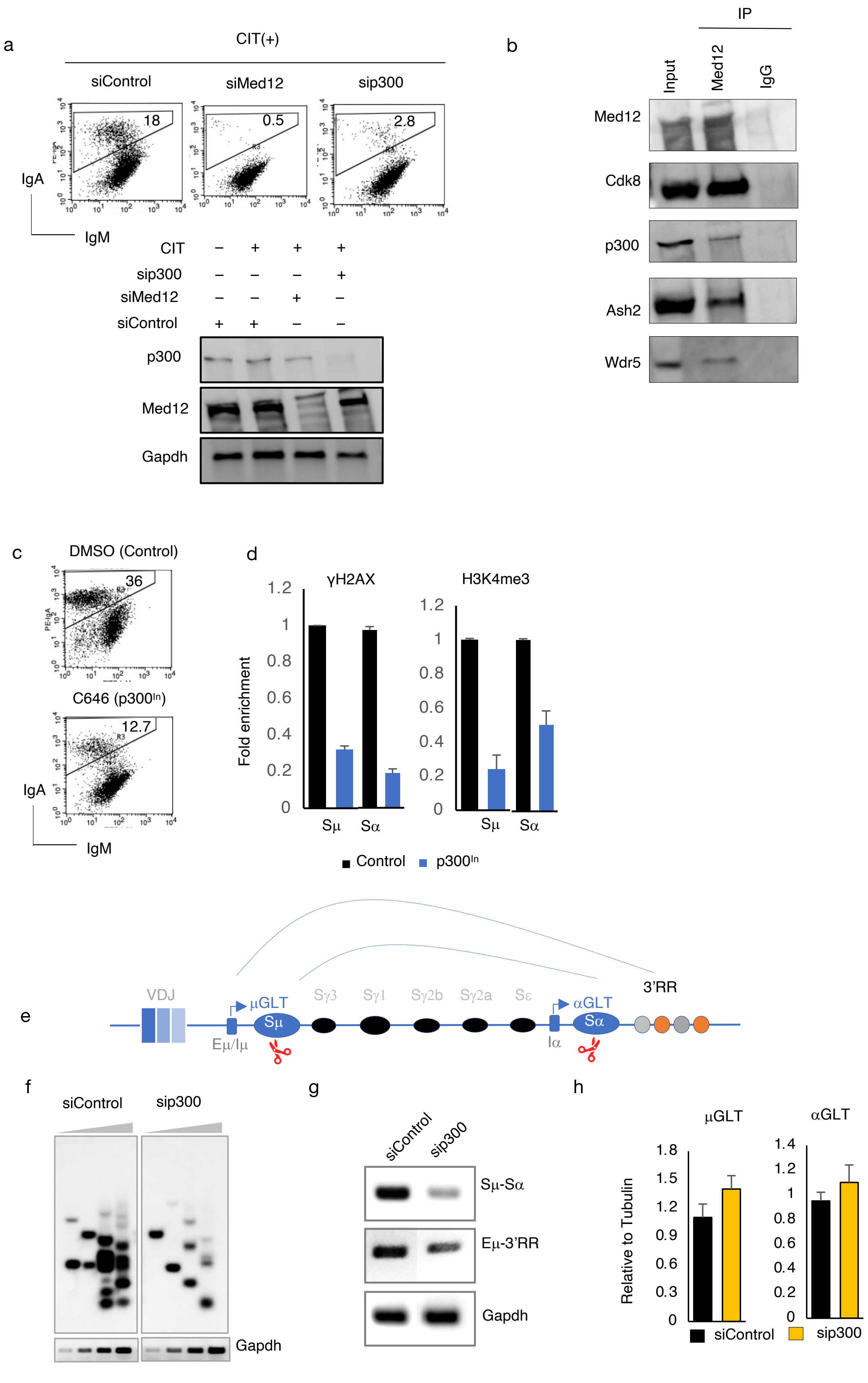

a
CIT(+)
siControl
siMed12
sip300
18
0.5
2.8
IgA
IgM
CIT
sip300
siMed12
siControl
p300
Med12
Gapdh
b
IP
Input
Med12
IgG
Med12
Cdk8
p300
Ash2
Wdr5
c
DMSO (Control)
36
C646 (p300In)
12.7
IgA
IgM
d
γH2AX
H3K4me3
Fold enrichment
Sμ
Sα
Control
p300In
e
VDJ
μGLT
Sμ
Sγ3
Sγ1
Sγ2b
Sγ2a
Sε
αGLT
Sα
3'RR
Eμ/Iμ
Iα
f
siControl
sip300
Gapdh
g
siControl
sip300
Sμ-Sα
Eμ-3'RR
Gapdh
h
μGLT
αGLT
Relative to Tubulin
siControl
sip300

**Extended Data Fig. 3**

**Med12-p300 complex is required for enhancer activation. a,** The siRNA against the Med12 and p300 showing their effect on CSR after 24h of CIT(+) in CH12F3-2A cells. The WB showing the confirmation of KD in each siRNA used. Gapdh were used as loading control.**b,** The WB blot showing the endogenous protein IP using Med12 or IgG control antibody. The blot was probed against the indicated antibodies. Med12 IP showing the interaction between p300 and FACT complex components. Cdk8 used as a positive control.**c**, The inhibition of p300 histone acetyltransferase by C646 inhibitor (HATi) and its effect on IgA switching. The stock solution of C646 was made in DMSO and the final 5mM was used as working concentration. DMSO without C646 was used as negative control. The IgA percentage was determined after 48h of CIT (+) stimulation.**d,** The effect of C646 inhibitor on AID induced DNA break formation. The ChIP-qPCR showing the relative enrichment of histone marks in DMSO control and C646 treated CH12F3-2A cells using indicated antibodies.**e,** Schematic scheme showing the long-range interaction between Switch and enhancers regions.**f**, LM-PCR showing the effect on $p300^{KD}$ on AID induced DNA breaks as described earlier. Gapdh used as internal control. The triangles indicate a threefold dilution in the DNA amount. **g,** 3C essay showing the long-range interaction between the indicated primer pairs in control and $p300^{KD}$ CH12-F3-2A cells. Gapdh PCR of the cross-linked DNA sample served as a loading control.**h,** The RT-qPCR showing the effect of control and $p300^{KD}$ on GLTs. The data was normalized to endogenous tubulin.

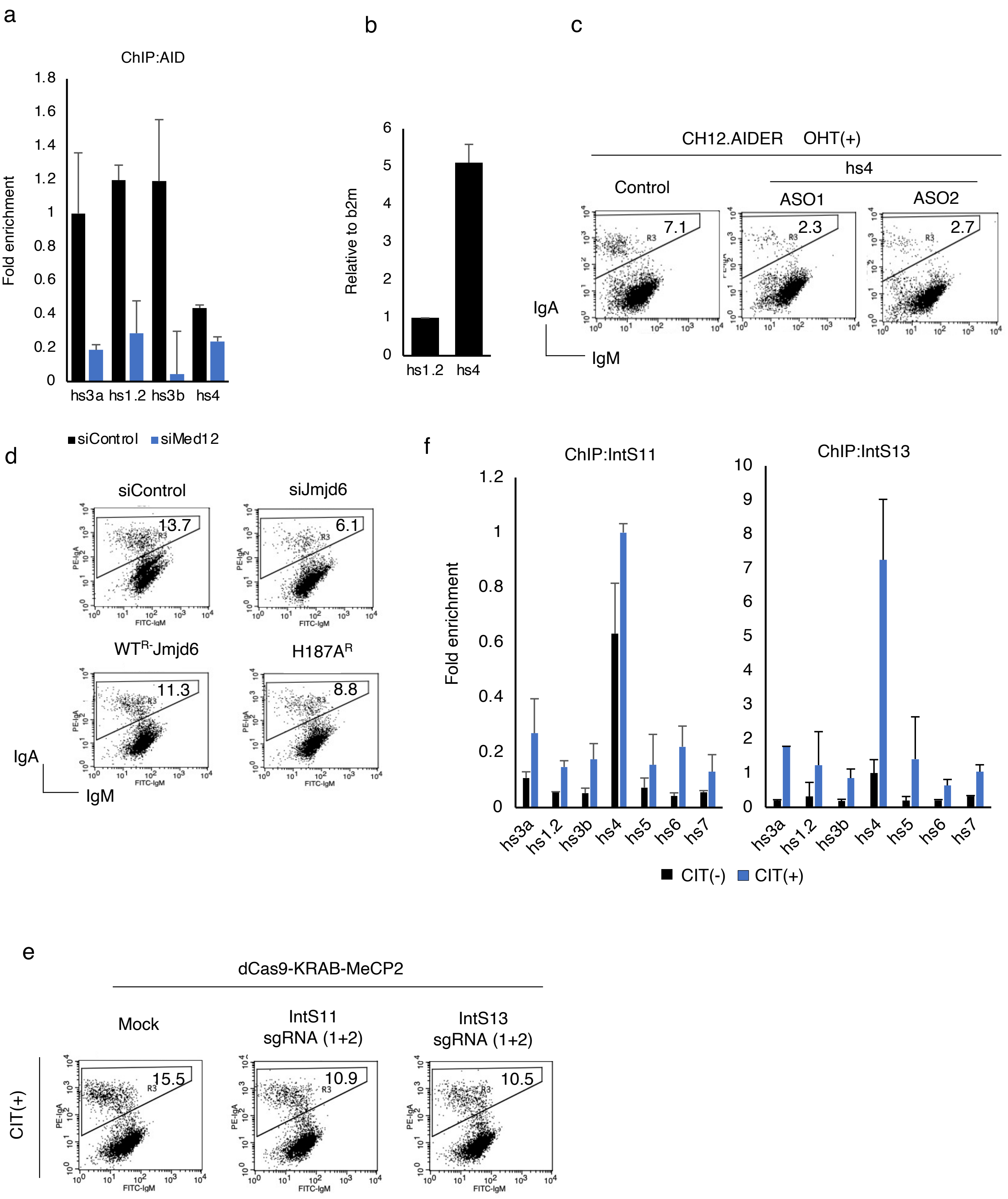
a
ChIP:AID
Fold enrichment
1.8
1.6
1.4
1.2
1
0.8
0.6
0.4
0.2
0
hs3a
hs1.2
hs3b
hs4
siControl
siMed12
b
Relative to b2m
6
5
4
3
2
1
0
hs1.2
hs4
c
CH12.AIDER OHT(+)
hs4
Control
ASO1
ASO2
7.1
2.3
2.7
R3
IgA
IgM
d
siControl
siJmjd6
13.7
6.1
WTR-Jmjd6
H187AR
11.3
8.8
PE-IgA
FITC-IgM
IgA
IgM
f
ChIP:IntS11
ChIP:IntS13
Fold enrichment
1.2
1
0.8
0.6
0.4
0.2
0
10
9
8
7
6
5
4
3
2
1
0
hs3a
hs1.2
hs3b
hs4
hs5
hs6
hs7
CIT(-)
CIT(+)
e
dCas9-KRAB-MeCP2
Mock
IntS11 sgRNA (1+2)
IntS13 sgRNA (1+2)
CIT(+)
15.5
10.9
10.5
PE-IgA
FITC-IgM

**Extended Data Fig. 4**

**Rescue of Med12 deficiency by CRISPR/dCas9 mediated 3'RR activation.a,** The AID ChIP qPCR showing the occupancy of AID at 3'RR regions in control and Med12 KD cells. The values were normalized to the DNA input signals followed by the maximum value in each data set. **b,** The RT-qPCR data showing the relative transcript level of hs1.2 and hs4 in unstimulated condition. The data were normalized with β2m.**c,**The FACS analysis showing the effect of hs4 specific ASOs on IgA efficiency after 24h of OHT (+) stimulation in AIDER cell line.**d,**The FACS analysis showing the effect of overexpressed $WT^{R}$-Jmjd6 and its catalytic defective mutant ($H187A^{R}$) on CSR in control and Jmjd6 depleted CH12F3-2A cells.**e,** The FACS analysis showing the effect of integrator gene knockdown on CSR using dCas9-KREB-MeCP2 targeted to integrator locus using 2 sets of pooled guide RNAs.**f,** The ChIP qPCR showing the occupancy of integrator components (Inst11/13) at 3'RR regions in CIT stimulated and unstimulated cells. The values were normalized to the DNA input signals followed by the maximum value in each data set.

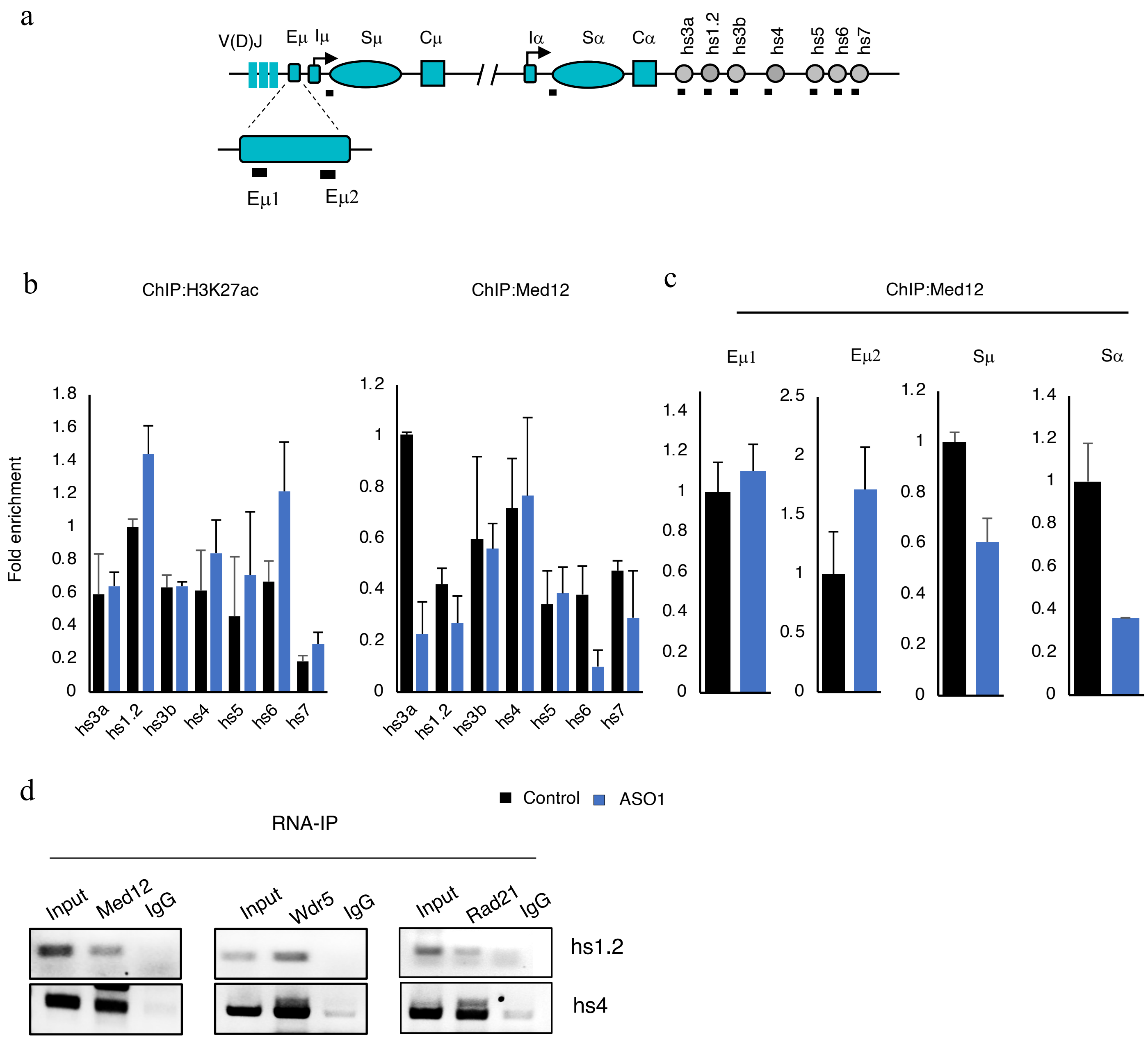


**Extended Data Fig. 5**

**Depletion of 3'RR eRNAs recruits Med12 at S region but not to enhancers via direct interaction.a,** The schematic showing the different regions present at the Igh locus, the black bar showing the position of the primers used for ChIP-qPCR amplification.**b,c,** The ChIP qPCR showing the enrichment in control samples and hs4 specific ASO1 knockdown CH12F3-2A cells using the indicated antibodies. The values were normalized to the DNA input signals followed by the maximum value in each data set.**d,** The gel picture showing the PCR performed after the RIP (RNA-IP) using the indicated antibodies. IgG served as a negative control.

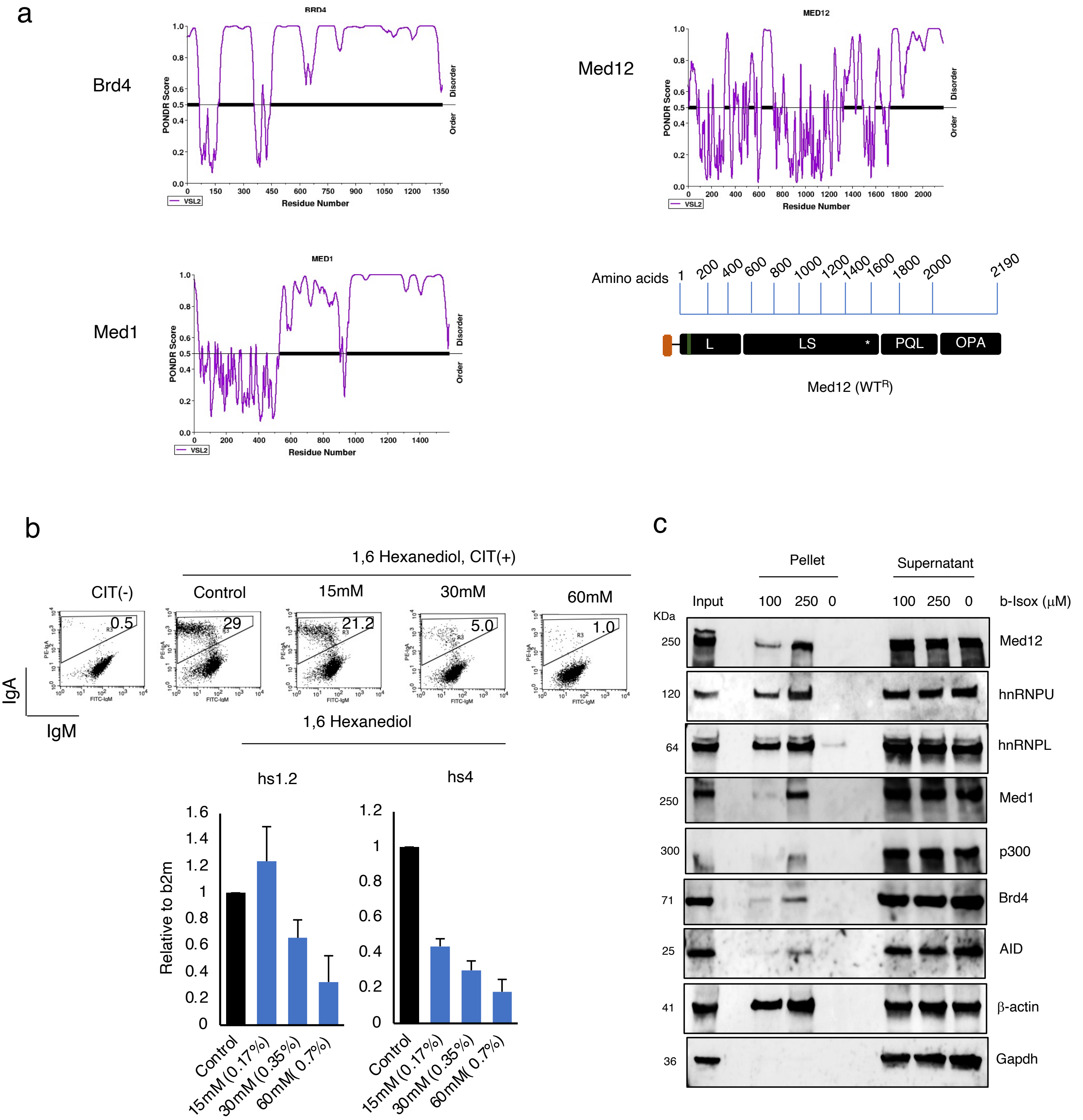


**Extended Data Fig. 6**

**The eRNA is required for DNA break-recombination specific condensates formation.a,**PONDR analysis showing the order/disordered regions present in indicated proteins.**b,** (Left) The FACS analysis showing the dose dependent effect of 1,6 hexanediol (1,6 HD) on CSR. (Right) The RT-qPCR analysis on different doses of 1,6 HD treated CH12F3-2A cells and its effect on hs1.2 and hs4 enhancer transcription. **c,** WB showing the biotinylated isoxazole (b-Isox) mediated low complexity protein precipitation involved in granule formations. Gapdh were used as negative control.

**Extended Data Table1**

**Med12 LS and PQL domains are crucial for CSR.** The table showing list of various disease associated Med12 mutants used in the study. Each color code represents the disease correspond to the respective domains and there effect on CSR.

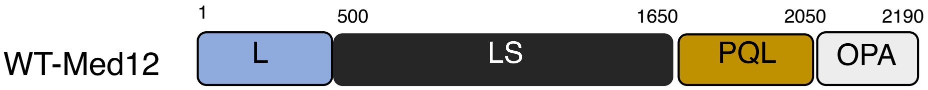


| Human Med12 | Mouse Med12 | Domain | Human Disease | CSR activity | Reference |
|---|---|---|---|---|---|
| WT | WT | | | 100 | |
| L36R | L36R | L domain | Uterine leiomyoma | 90±8 | (1) |
| Q43P | Q43P | L domain | Uterine leiomyoma | 92±7 | (1) |
| G44S | G44S | L domain | Uterine leiomyoma | 85±15 | (1) |
| E172Q | E172Q | L domain | OSMKB | 75±18 | (2) |
| R296E | R296E | L domain | Phenotypic abnormality | 88±10 | (2) |
| R521H | R520H | LS domain | XLID-atypical | 54±7 | (3) |
| R621Q | R622Q | LS domain | XLID-atypical | 65±4 | (4) |
| D727E | D728E | LS domain | Prostrate cancer | 90±10 | (5) |
| I771T | I772T | LS domain | XLID-atypical | 60±10 | (6) |
| N898D | N899D | LS domain | Similar to FG | 87±7 | (7) |
| G958E | G959E | LS domain | XLID-FG syndrome | 76±10 | (8) |
| R961W | R962W | LS domain | XLID-FG syndrome | 75±9 | (9) |
| N1007S | N1007S | LS domain | XLID-Lujan Syndrome | 60±8 | (10) |
| R1138H | R1139H | LS domain | This study | 42±7 | This study |
| R1148H | R1149H | LS domain | XLID-Ohdo syndrome | 93±4 | (11) |
| S1165P | S1166P | LS domain | XLID-Ohdo syndrome | 90±8 | (11) |
| E1091K | E1092K | LS domain | Prostrate cancer | 78±14 | (2) |
| L1224F | L1225F | LS domain | Prostrate cancer | 58±5 | (12) |
| R1295H | R1296H | LS domain | Disturb IEG expression | 92±7 | (7) |
| P1310Q | P1311Q | LS domain | Prostrate cancer | 75±22 | (5) |
| A1383T | A1384T | LS domain | XLID-Ohdo syndrome | 90±3 | (13) |
| R1463C | R1462C | LS domain | This study | 40±8 | This study |
| H1729N | H1731N | PQL domain | XLID-Ohdo syndrome | 56±10 | (11) |
| N1845T | N1847T | PQL domain | Prostrate cancer | 80±10 | (5) |
| R1862K | R1864K | PQL domain | Methylation defective | 43±8 | (14,15) |
| R1899K | R1901K | PQL domain | Methylation defective | 50±4 | (14) |
| R1912K | R1914K | PQL domain | Methylation defective | 90±10 | (14,15) |

| Antibody | Company | Catalouge Number |
|---|---|---|
| **Western Blot antibodies** | | |
| Anti-AID | Cell singanalling technology | 30F12 |
| Anti- alpha Tubulin | Merck | CP06 |
| Anti-Flag | Sigma | F3165 |
| Anti-Med12 | abcam | ab70842 |
| Anti-CDK8 | Cell singanalling technology | 17395 |
| Anti-EP300/KAT3B | abcam | ab275378 |
| Anti-Ash2l | Millipore | ABE1972 |
| Anti-WDR5 | Cell singanalling technology | 13105 |
| Anti-Top1 | Santa Cruz | sc-5342 |
| Anti-hnRNPU | abcam | ab10297 |
| Anti-hnRNPL | Santa Cruz | sc-48391 |
| Anti-B-actin | Sigma | A1978-200ul |
| Anti-Gapdh | Milipore | MAB374 |
| Anti-Med1 | Milipore | 17-10530 |
| Anti-Brd4 | abcam | ab75898 |
| IgM-FITC | eBiosciences | E00715-1631 |
| IgA-PE | Southern Biotech | C2904-W100 |
| **ChIP/RIP antibodies** | | |
| Anti-Med12 (ChIP) | Bethyl | A300-774A |
| Anti-Jmjd6 (ChIP) | Biorbyt | orb340986 |
| Anti-H3k27ac(ChIP) | Active motif | 39133 |
| Anti-Carm1 (ChIP) | Cell singanalling technology | 12495 |
| Anti-Ints11 (ChIP) | Bethyl | A301-274A |
| Anti-Ints13 (ChIP) | Bethyl | A303-575A |
| Anti-Rad21 (RIP) | abcam | ab992 |
| Anti-gH2AX (ChIP) | Miilipore | 05-636 |
| Anti-H3K4me3 (ChIP) | Millipore | 07-473 |
| Anti-IgG (ChIP) | Milipore | PP64B |
| Anti-AID (ChIP) | Invtirogen | 39-2500 (ZA001) |
| Anti-P300/KAT3B (ChIP) | abcam | ab275378 |
| Anti-Med12 (RIP) | abcam | ab70842 |
| Anti-Wdr5 (RIP) | Cell singanalling technology | 13105 |

| List of siRNA used | | | Sequence |
|---|---|---|---|
| mhs4-500_1 ASO | Qiagen | 339511 LG00788451-DDA | TGCTCAGATGGATTGC |
| mhs4-500_2 ASO | Qiagen | 339511 LG00788452-DDA | TCAGGATCCACAACTA |
| mhs4-500_4 ASO | Qiagen | 339511 LG00788454-DDA | ACCAGCCAAGACAATA |
| siMed12 | Ambion | s81804 | CCAUCUACUGUAACGUGGA |
| siJmjd6 | Invitrogen | MSS272577 | CAGGAUGAAGCAAUUACCUGGUUUA |
| siMed13-1 | Invitrogen | MSS220711 | CCUACAUCUCCUGAGGAAGUCCAAA |
| siMed13-2 | Invitrogen | MSS283064 | GAAGCUAAUGUAGCCUGGAAGUAUU |
| siMed13-3 | Invitrogen | MSS220710 | GAGGAGAAACAGGACGAUAUGGAUU |
| siMed13L-1 | Invitrogen | MSS233466 | GAGGAGCUUGUGAGCGAGUGGUUUA |
| siMed13L-2 | Invitrogen | MSS233467 | UCCCUUUAUAUUGGCUCCAAUCAAA |
| siMed13L-3 | Invitrogen | MSS293541 | CCGCCAUCACCUAGCACCUUAUUUA |
| siCdk8-1 | Invitrogen | MSS281888 | CAGCUGGACAGAAUAUUCAAUGUAA |
| siCdk8-2 | Invitrogen | MSS281886 | CGAAAGCCAACAAGAAGCCAGUUCA |
| siCdk8-2 | Invitrogen | MSS281887 | GACAUGGGCUUUGCCCGAUUAUUUA |
| sihMed12-1 | Ambion | HSS115114 | GGGACAUUGGCUGCAGACAUGUCUA |
| sihMed12-2 | Ambion | HSS115115 | GCACCACCCAAACCCUGGUUCUAUA |
| sihMed12-3 | Ambion | HSS190751 | CGACUCCCUUUAGUGACCUGCUUAU |
| siCarm1 | Invitrogen | MSS226868 | CCAACUUCUGGUACCAGCCAUCCUU |
| siP300 | Invitrogen | MSS220767 | CCUGGACAGUCAAAGAAGAAGAUUU |

| Construct | Site Directed Mutagenesis Primers | |
|---|---|---|
| | (Forward 5'-3') | (Reverse 3'-5') |
| pCMV6-3XFlag-WT-mMed12$^{R}$ | ATATATCGTGTTCTTCACTTTTGAGCAAAAGTCACTGAAGAGCTC | TGCAATGTAGAACCATCGGAATCCAATATGCGCTGG |
| pCMV6-3XFlag-ΔL-mMed12$^{R}$ | GATGATGATGCTGTGGTATCATTATTGTGTGAATGGGCT | GGGAGGCCCCAGCCGCAGCCGCTTCAGG |
| pCMV6-3XFlag-ΔLS-mMed12$^{R}$ | AAGCTTCAGAAGGACTTGGGGGAGCGCCAATCAGA | CAATAATGATACCACAGCATCATCATCAGAGGAAA |
| pCMV6-3XFlag-ΔPQL-mMed12$^{R}$ | GGCGTCCGTTCAACTTCCATCCTGCCTGAG | CTTCACCAGGTTCATATATGCACGCTTGTTTTCCTCCATGCT |
| pCMV6-3XFlag-ΔOPA-mMed12$^{R}$ | GCTTTGGTCCGGCAACTTCAACAACAGCTCTCTAATACC | ACCCATCATGGGTGTCTGCTGCAGTGTCTGGTGTGAAAA |
| pCMV6-3XFlag-ΔNLS-mMed12$^{R}$ | CCTCCCGATGTGTACCCTCAAGATCCCAAA | TTCATAGCTCAAGATCCCGAAAGCCGCCAT |
| pCMV6-3XFlag-L36R-mMed12$^{R}$ | AACAGAAGGAGGATGAAAGGACGGCTTTGA | TGGGATCTTGAGGGTACACATCGGGAGG |
| pCMV6-3XFlag-Q43P-mMed12$^{R}$ | TTGAATGTAAAACCAGGTTTCAATAACCAGCCTG | AGCCGTCAGTTCATCCTCCTTCTGTTTGGG |
| pCMV6-3XFlag-G44S-mMed12$^{R}$ | GGCTTTGAATGTAAAACAAAGTTTCAATAACCAGCCT | GTCAGTTCATCCTCCTTCTGTTTGGGATCTTG |
| pCMV6-3XFlag-E172Q-mMed12$^{R}$ | GAAAAATACTGCTGACCCCTTCACTGAATGGAC | TTCTTAACCTTAGTCTGAGACATTGCTGCATAGTAGG |
| pCMV6-3XFlag-R296E-mMed12$^{R}$ | TACCCGGAGATTGGCTCTCCAGCTGG | CAGAAGTAGGCAAGTTCGCGGGACAGATAGGCA |
| pCMV6-3XFlag-R520H-mMed12$^{R}$ | CCAAACTCCTAGAGAAGAGACAAGCAGAAA | CTACTACCATAGCATGATGCCGACCTGAGCGTT |
| pCMV6-3XFlag-R622Q-mMed12$^{R}$ | CCCTGGTCCTCGGCCTCCCTCTCCCTT | GCTCCAAATGCAAGATCCCCCTGAGAGATGAGAGTGC |
| pCMV6-3XFlag-D728E-mMed12$^{R}$ | CAGTATGCCACACACTTTCCAATCCCACAG | CACATGTCGTGGCTGCTCATACAGAATCCCAAGTG |
| pCMV6-3XFlag-I772T-mMed12$^{R}$ | GGATATCCTGAAGGTTCTGAATCGCAAGGGGACA | TTGGTAATCTTCTTGGTGGCATGGCGGGCATCA |
| pCMV6-3XFlag-N899D-mMed12$^{R}$ | GGCGGAGCTCCTTCTCAAATCGTCGGATCTGG | TCAACCACACTCAGCTCGTCCAGTAACTGGATGGCAAAG |
| pCMV6-3XFlag-G959E-mMed12$^{R}$ | ATGGCTCCTCTGCAGAGCGCTGTATC | CTGAACGGTTCATTTCATGTTTCACCACGCCAC |
| pCMV6-3XFlag-G962W-mMed12$^{R}$ | AAACATGGAATGAACTGGTCAGATGGCTCCTCTG | CACCACGCCACAGAGCCCCTCAAA |
| pCMV6-3XFlag-N1007S-mMed12$^{R}$ | ACACGATATATTGCAGCGTAGAACCATCGGAAT | TCTTCACTTTTGAGCAAAAGTCACTGAAGAGC |
| pCMV6-3XFlag-R1139H-mMed12$^{R}$ | AAGACCTGATTCGCTGTGCAGCCATCCC | CTAGGAGCAAACATTGACGGGCGATGAGGATAGC |
| pCMV6-3XFlag-R1149H-mMed12$^{R}$ | TAGAAGACCTGATTCACTGTGCAGCCATCCC | GGAGCAAACATTGACGGGCGATGAGGATAGC |
| pCMV6-3XFlag-S1166P-mMed12$^{R}$ | TAGTGAGCAGGATCCTGAGCCAGGAGCCAG | CAAGCAGCATTGAGGAGTGAAGGGATGGCTGCAC |
| pCMV6-3XFlag-E1092K-mMed12$^{R}$ | TAAGGCCTTGTGCTGCTCCTCGAACAATGGCAC | AGTACTCCTAGCCACTTAGCACTCAAGGACTTGCAA |
| pCMV6-3XFlag-L1225F-mMed12$^{R}$ | ACTCGGAGATGCGGAGCTAAAAGGTTCAGGC | ACAAACACAGCCTTGAAAACAGCAAACACAGC |
| pCMV6-3XFlag-R1296H-mMed12$^{R}$ | GTGTGAGGATAGCAATGATCTACAAGACCCAGTG | AGTGACTTAAGGCAATGTTCTCCTACCCATTCCTGTTGG |
| pCMV6-3XFlag-P1311Q-mMed12$^{R}$ | GCCCAGCGCCTCATGCAACTTATCTGCTAC | CTGGGCACTACTCAACACTTGGTCTTGTAGATCATTGCT |
| pCMV6-3XFlag-A1384T-mMed12$^{R}$ | CTTGGAGAACATTACCAAGGCCACAATCGAGGTTTT | CTAGGAGCAAACATTGATGGGCGATGAGGATAGCAAC |
| pCMV6-3XFlag-R1462C-mMed12$^{R}$ | CAGCACCTGGGCTCTTCTTCCTGTAAAGAACGAGATCGACAGAAA | ACCCTTTTCCAGTTCCTCCCCAGCAGCTTTTAATACATGGCC |
| pCMV6-3XFlag-H1731N-mMed12$^{R}$ | TTGCTCTATCATACCAACCTGAGGCCTCGAC | CAGCCGCTGCTGCTCCTCCCCTCGTG |
| pCMV6-3XFlag-N1847T-mMed12$^{R}$ | CTGGTGGCCCTCGTGTGGATCCATACCGC | CAGGTAGTGGCTGGGTTTGGGTATACAGGCCCA |
| pCMV6-3XFlag-R1864K-mMed12$^{R}$ | CGCCCCGTGAAATTACCAATGCAAAAGCTGC | GTATGGATCCACACGAGGGCCACCAGCAG |
| pCMV6-3XFlag-R1901K-mMed12$^{R}$ | TCTGTATACAAGCAGCAGCAACCCACAGTGCCCCA | TGTCTTATAAGAAGAGGGTTCTAGGCCCATGACAGTA |
| pCMV6-3XFlag-R1914K-mMed12$^{R}$ | CAACAGCTCCAGGCAAAGATACAGAGTCAGGG | TTTAAGGCGCTGTCCCTGGGGCACTGT |
| pCMV6-3XFlag-WT-mJmjd6$^{R}$ | ATAACGTGGTTCAATGTCATCTATCCCCGGACACA | ATAACGTGGTTCAATGTCATCTATCCCCGGACACA |
| pCMV10-3XFlag-H187A-mJmjd6$^{R}$ | GGCAATCCCAGTTCCAGAACGCGGTGG | ATCGACCCTCTGGGGACCAGTGCCTG |

| qPCR primers | (Forward 5'-3') | (Reverse 3'-5') |
|---|---|---|
| μGLT | CTCTGGCCCTGCTTATTGTTG | AATGGTGCTGGGCAGGAAGT |
| αGLT | CCAGGCATGGTTGAGATAGAGATAG | GAGCTGGTGGGAGTGTCAGTG |
| mAID | CGTGGTGAAGAGGAGAGATAGTG | CAGTCTGAGATGTAGCGTAGGAA |
| Med12 | GAAGTACAGCAAGCAACATGCCCAGCAG | CAATCTGGTGCACCTGGCTGTGGAGGG |
| Egr1 | ATCGCTCTGAATAATGAGAAGGCGAT | CACTGACTAGGCTGAAAAGGGGTTCA |
| Atf1 | ATCTGACAGCATAGGCTCCTCACAGA | TTGGGGCAATGGCAATGTACTGTCC |
| Atf7 | GCCTCTTGACATGTCTCTGCCT | CAGTGGTGGGGAACAGGGG |
| Cdk8 | ACTACTAACGTCAGAACCAATATTTCACTGT | CTTTCATTAATGTTGAATGTTCGGGCATCTT |
| Med13 | GAGCTTCTCACTGTTCAGCTTCTTGC | GCTAGTACTGTCAGTGCTGAAGCCAT |
| Med13L | GACAGCACAAGCCCATCTGCAAAGT | ATGGCGGCAAACTTGAGCATAAAGT |
| hs1.2 | CCCCAGATTGTAACCCTAGCC | TCTCCATGATGGCCTAGCCT |
| hs4 | AGGAAGTCCTCTGTGACTCG | ACTTCCATCAGGTCAACCTCC |
| P300 | GGATTAAGTTTGATAAATAGCATGGTCAAA | ACCTGCTGGGGACTGAGTAGGA |
| Jmjd6 | ACAAACACACCCAGAGAACTCATCAAG | CCAGGGGTTTGAATTCAGGTGGC |
| β2M | ACAGTTCCACCCGCCTCACA | TTGCTGAAGGACATATCTGACATCTCTACT |
| Carm1 | AGAGTAACAATCTGACAGACCGCATCGTGG | ACTTTTTGGCATGGAGGTAGCTCTCGAGC |
| Tubulin | TTCAAGGTTGGCATCAACTACCAGC | TGGCTGTGGTGTTGCTCAGCATGC |

| LM-PCR | | |
|---|---|---|
| LMPCR.1 | GCGGTGACCCGGGAGATCTGAATTC | |
| LMPCR.2 | GAATTCAGATC | |
| 5′ Sμ | GCAGAAAATTTAGATAAAATGGATACCTCAGTGG | GCGGTGACCCGGGAGATCTGAATTC |
| 5′ Sμ-probe | DIG-AGGGACCCAGGCTAAGAAGGCAAT | |
| GAPDH | ATCCTGTAGGCCAGGTGATG | AGGCTCAAGGGCTTTTAAGG |

| 3C essay | | |
|---|---|---|
| Eμ | GGAACAATTCCACACAAAGACTC | |
| Eα (3'RR) | CAAGGTGTTAAGGAAAACTTGCTC | |
| Sμ | GCTGACATGGATTATGTGAGG | |
| Sα | GAGCTAGGCTAGACTTACTAAGC | |
| GAPDH3c | AGTAGTGCGTTCTGTAGATTCC | CAGTAGACTCCACGACATAC |

| ChIP-qPCR primers | | |
|---|---|---|
| Eμ1 | GGAGTGGGGCACTTTCTTTAGA | CAGCAACTACCCTTTTGAGACCG |
| Eμ2 | CAGGCACCGCAAATGGTAAG | TGTGAGTGACCCAGACAACG |
| Iμ | AAGGGCTTCTAAGCCAGTCC | CACAACCATACATTCCCAGGT |
| Sμ | CAATGTGGTTTAATGAATTTGAAGTTGCCA | TCTCACACTCACCTTGGATCTAAGCACTGT |
| Cμ | CAGCACCATTTCCTTCACCTGGAACTACCA | GGCTAGGTACTTGCCCCCTGTCCTCAGTGT |
| Iα | GTGATTCAGGGAGCAAGAGC | TCTAGCCTGGGAGTCTCCTG |
| Sα | TGAAAAGACTTTGGATGAAATGTGAACCAA | GATACTAGGTTGCATGGCTCCATTCACACA |
| Cα | AGTGCCCAGAAGGAGAATCCGTGAAATGTT | GACCCCCTAACGTTCTTTACCAGAGCAATT |
| hs3a | GGGTAGGGCAGGGATGCTCACAT | GCTATGATCAAAACCAGTCATCGG |
| hs1.2 | AGCATAGGCCACTGGGACTGG | CTCTCACTTCCCTGGGGTGTT |
| hs3b | CTGACAGAGACAGACAGCGTTCC | TGGTTTGGGGCACCTGTGCTGAG |
| hs4 | GGGTAGGGCAGGGATGTTCACAT | CCATGGGACTGAAACTCAGGGAACCAGAAC |
| hs5 | CTCTGTGACTCGTCCTTAGG | GCTTGCCTTCTGGGCTAGAAC |
| hs6 | AGCAGCCAGTGCTCTTAACCC | GTTGAGTCTGAAGATACACGCG |
| hs7 | TCGCAGCTATTCCCAACTACG | AATACGTTCCTAGGGCCTAACGATACT |

| sgRNA oligos for CRISPRa | 5'-3' |
|---|---|
| hs1.2 | GTCTGCACAGAATGAGTAACT |
| hs4 | GCTCGGGGAACCAGAACCGTA |
| **sgRNA oligos for CRISPRi** | |
| IntS11 (1) | TGGGTGAGTCCGAGTCCTA |
| IntS11 (2) | TGGAGCCGTGGTTCGAGGCG |
| IntS13 (1) | GCCGTGGCTTAGTACTTTGT |
| IntS13 (2) | AGTGCGCTCTTGTGCTTGCA |

| Translocation | |
|---|---|
| Su5'1a(1st) | ACTATGCTATGGACTACTGGGGTCAAG |
| c-myc5′1a(1st) | GTGAAAACCGACTGTGGCCCTGGAA |
| Su5'1b(2nd) | CCTCAGTCACCGTCTCCTCAGGTA |
| c-myc5′1b(2nd) | GTGGAGGTGTATGGGGTGTAGAC |
| *myc* probe | GGACTGCGCAGGGAGACCTACAGGGG |
| *Igh* probe | GAGGGAGCCGGCTGAGAGAAGTTGGG |
